# Double-edged role of resource competition in gene expression noise and control

**DOI:** 10.1101/2021.10.29.466467

**Authors:** Hanah Goetz, Austin Stone, Rong Zhang, Ying-Cheng Lai, Xiao-Jun Tian

**Author notes:** These authors contributed equally to this work (Hanah Goetz, Austin Stone).

## Abstract

Despite extensive investigation demonstrating that resource competition can significantly alter the circuits’ deterministic behaviors, a fundamental issue is how resource competition contributes to the gene expression noise and how the noise can be controlled. Utilizing a two-gene circuit as a prototypical system, we uncover a surprising double-edged role of resource competition in gene expression noise: the competition decreases noise through a resource constraint but generates its own type of noise which we name as “resource competitive noise.” Utilization of orthogonal resources enables retaining the noise reduction conferred by resource constraint while removing the added resource competitive noise. The noise reduction effects are studied using three negative feedback controller types: negatively competitive regulation (NCR), local, and global controllers, each having four placement architectures in the protein biosynthesis pathway (mRNA or protein inhibition on transcription or translation). Our results show that both local and NCR controllers with mRNA-mediated inhibition are efficacious at reducing noise, with NCR controllers demonstrating a superior noise-reduction capability. We also find that combining negative feedback controllers with orthogonal resources can improve the local controllers. This work provides deep insights into the origin of stochasticity in gene circuits with resource competition and guidance for developing effective noise control strategies.

## INTRODUCTION

The last three decades have witnessed an increasing exploitation of synthetic gene circuits in clinical applications for diagnostics and therapeutics. In a gene circuitry, the competition over transcriptional and translational resources between multiple modules is essential and has been demonstrated to play a significant role in regulating the deterministic behaviors of the synthetic circuits [1–3]. For example, how resource competition can alter the means of circuit species and even completely change the steady states of the dynamical systems has been studied [4–6]. While gene circuits can exhibit certain deterministic behaviors to some extent, they are intrinsically stochastic, and this can significantly affect the circuit function. In fact, noisy protein expression is a common mode for circuit malfunction and design failure. In view of the importance of resource competition in the circuit dynamics, it is of fundamental interest to investigate how the competition affects or contributes to the stochastic nature of the circuits. To understand the origin of stochasticity in gene circuits is important not only for better understanding the dynamics but also for advancing gene circuit engineering.

A closely related issue is control. In general, the effects of resource competition can be attenuated in orthogonal resource systems and by negative feedback control. For example, orthogonal ribosomes and RNA polymerases (RNAPs) have been engineered for creating separate resource pools for genes [7–11], and various circuit controller topologies have been exploited to mitigate the burden that the components have on each other without affecting the overall function of the circuit [12–18]. The existing controllers utilize some sort of negative feedback topology to repress circuit outputs when the synthetic circuit begins taking up more than its fair share of the host’s resources. Three categories of such controllers have been studied: global [19], local [15], and negatively competitive regulatory (NCR) controllers [18] controllers. While these control strategies have been demonstrated to reduce resource competitive effects, it is unclear whether they can reduce gene expression noise and, if yes, which architecture represents the optimal design for noise control.

Given the universal existence of resource competition in gene circuits and considering that the traditional protein expression models do not take into consideration resource constraints, it is of fundamental importance to uncover and understand the competition-induced noise and characterize the stochastic behavior for synthetic biological constructs. As feedback control serves to reduce noise, it is also imperative to evaluate the efficacy of different control strategies to identify the optimal one. This paper addresses these issues. First, we analytically compare the noise in the idealized scenario in which resources are unlimited with the realistic case subject to the constraint of resource competition. This analysis leads to a new type of noise derived from resource competition and a reduction in noise caused by the resource limitation constraints, revealing a striking double-edged effect of resource competition on the noise behavior. We then analyze how the addition of orthogonal resources can remove the competition-induced noise. Finally, we compare the noise reduction performance of three general types of negative feedback controllers: NCR, local, and global, as well as each of their four placement subtypes. A useful finding is that global controllers are not effective at reducing noise and often can even increase noise, but combining negative feedback controllers with orthogonal resources can improve the local controllers and make the transcriptional inhibition strategies more effective. Our results provide unprecedented insights into the origin of stochasticity in synthetic gene circuits as well as into developing effective noise control strategies for resource competitive systems.

## RESULTS

### Double-edged effects of resource competition on gene expression noise

We consider a circuit with two independently regulated genes in the same cell (GFP and RFP), as shown in Fig. 1a, which is a prototypical design for characterizing gene expression noise [20] and resource competition [4, 17]. While the competition is not significant with only one copy of the two genes [20], it can become significant if high-copy plasmids are present. As in models for gene expression noise [21–28], we consider the transcription for the production of gfp and rfp mRNAs, the translation for the production of proteins GFP and RFP, and the degradation of both mRNAs and proteins. A feature of the existing models is that the transcription rate per gene and the translation rate per mRNA are constants, which is idealized. Going beyond the existing studies, we incorporate the competition for shared transcriptional and translational resources between the two genes into our model, which makes the transcription and translation fluctuate dynamically as in real gene circuits. The dynamical fluctuations are effectively random, leading to changes in the stochastic behaviors of the system. Resource competition between multiple genetic nodes in the same cell can have significant effects on the deterministic behaviors of the circuit. For example, the sharing of resources can create indirect inhibition links between the genes (indicated by the dashed lines in Fig. 1b) because, when one genetic module pulls from the resource pool, there are fewer resources available to the other module.

**FIG. 1.**
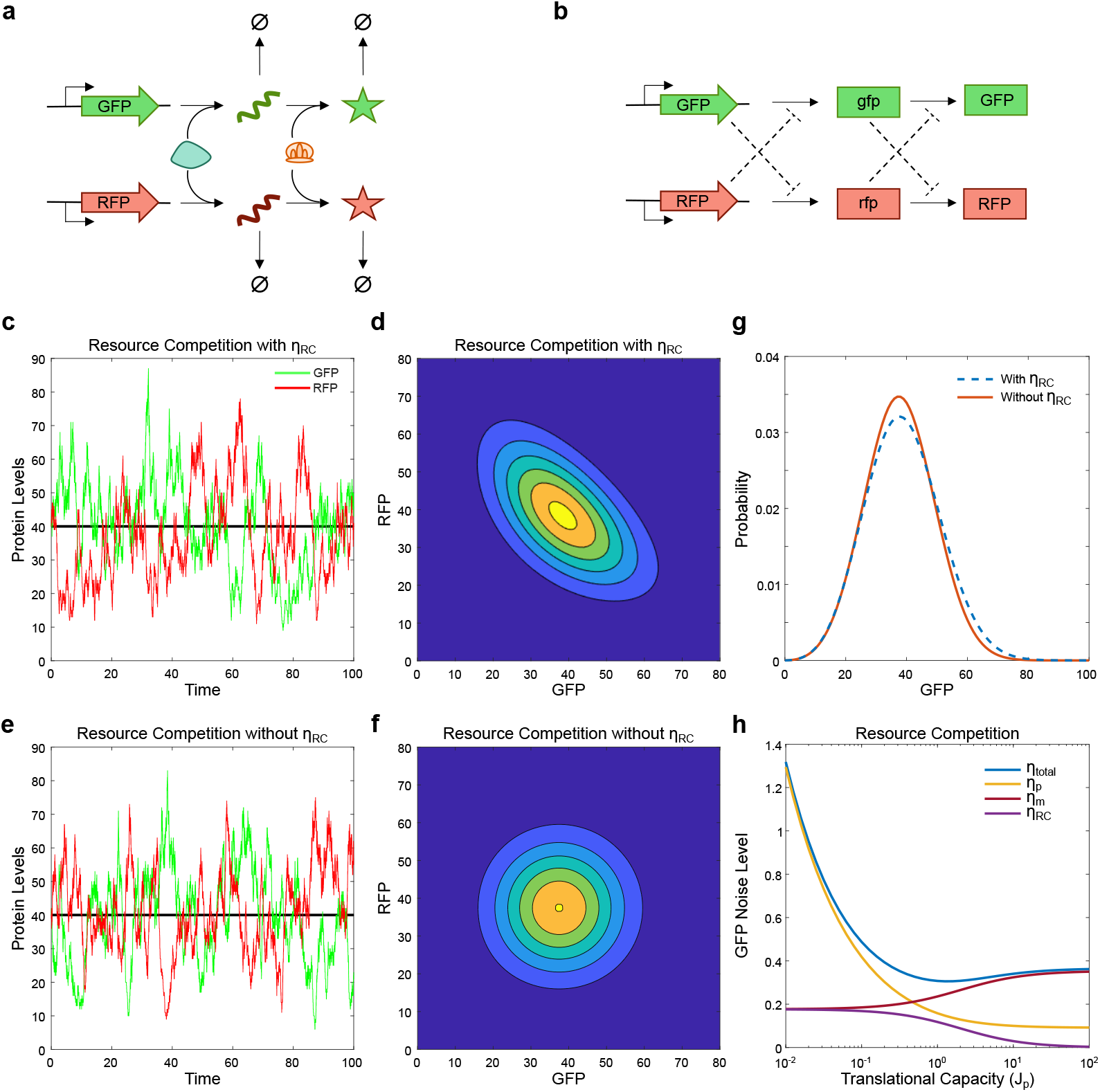
Double-edged effects of resource competition on gene expression noise. (a) Schematic diagram of GFP and RFP expression through shared transcriptional resources (RNAPs) to create mRNAs and translational resources (ribosomes) to create proteins. (b) Diagram illustrating how resource competition creates inhibition between the circuit modules. (c) Gillespie stochastic trajectories of GFP (green trace) and RFP (red trace) expressions. (d) Distribution of GFP and RFP expression levels obtained by solving the master equation in the two-gene circuit coupled by resource competition. (e) Gillespie stochastic trajectories of GFP (green) and RFP (red) expression. (f) Distribution of GFP and RFP expression levels in the two-gene circuit, in which the contribution of the fluctuation of one mRNA to the noise of the other protein due to resource competition has been removed. The horizontal black lines in (c) and (e) indicate the mean of proteins. (g) Distribution of GFP expression levels in the case with resource competition noise (blue dashed curve) and with the contribution of the fluctuation of one mRNA to the noise of the other protein due to resource competition removed (solid red curve). (h) FDT analytical solutions of the dependence of the total protein noise (blue curve), noise from the stochastic birth/death of protein (yellow curve), noise from the fluctuation of its own mRNA (maroon curve), and noise from the other mRNA (purple curve) on the translational capacity *J*_*p*_ of limited resources in the host cell for the synthetic gene circuit.

We begin by constructing two mathematical models for the two-gene circuit, one with unlimited resources (without competition) and another with competition, with details given in Method A. In the model with unlimited resources (UR model), the translation rate in each module depends linearly on the concentration of its own mRNA, as in previous models [21, 23, 24, 26–28]. In the model with resource competition (RC model), the translation rate depends on the concentrations of all mRNAs in the system [6, 18]. For simplicity, we do not include extrinsic noise from fluctuations in other cellular components of the system. To fairly compare the stochastic behaviors of the RC and UR models, we rescale the transcription and translation rate constants in both models to ensure that they have the same mean values of the mRNAs and proteins. We use the standard Gillespie method [29] (Method B) to generate stochastic trajectories, as shown in Fig. 1c, where the peaks in the GFP level more often correspond with valleys in the RFP level in the RC case. Likewise, peaks in RFP correspond with valleys in GFP. That is, the two proteins fluctuate in an anticorrelated fashion. This is an excellent manifestation of inhibition at work. A rise in one protein expression implies that its corresponding gene has used more resources. As a result, there are fewer resources available for the opposing gene, leading to a decrease in its protein expression. This anticorrelation is confirmed with the 2D GFP/RFP probability distribution obtained as a solution to the system’s master equation (Method C), as shown in Fig. 1d, where the deep blue color represents regions of low probability, while yellow represents the regions of the highest probability. It can be seen that a high GFP level mostly corresponds to low RFP levels and vice versa.

In the UR model, stochastic trajectories and their probability distribution indicate that expressed GFP is not related to the expression of RFP, as shown in Figs. 1a and 1b. That is, the two genes remain completely unconnected given that the inhibitions from resource competition are not present. We compare the expression distribution of GFP under conditions with unlimited and limited resources and find that the resource competition narrows the GFP distribution, as shown in Fig. 1c. This result implies that resource limitation provides the benefit of noise reduction even though it also leads to anticorrelation between the expression of two genes when compared to a case of unlimited resources.

From simulations, we notice that, due to resource competition, the fluctuations of one mRNA can contribute to the noise of the other protein. This contribution can be eliminated by setting the concentration of one mRNA to a constant (e.g., its mean) in the production rate of the other protein, thereby keeping the fluctuations of one gene’s expression from being a factor in the opposing gene’s expression. By so doing, we are able to determine the contribution of the resource competition to the total noise. Once eliminated, the anticorrelation disappears. That is, protein expression peaks in the stochastic trajectory can align with either the opposing protein’s peaks or the valleys, as shown in Fig. 1e. The 2D GFP/RFP probability distribution now becomes circular, as shown in Fig. 1f, in contrast to the ovular in Fig. 1d. It can be seen that high expression areas are now confined to a smaller region of the phase space than in the original RC case. This difference can also be seen in the 1D GFP probability distribution, as shown in Fig. 1g, where the distribution in the RC case (dashed blue curve) is lower and wider than that in the case where the fluctuations of one gene’s expression to the other have been eliminated (solid red curve). The results in Figs. 1c-1g thus indicate that the extra noise included in the RC case can be attributed entirely to resource competition. We call this “resource competitive noise,” denoted as *η*_*RC*_.

To better see the double-edged effects of resource limitation on the noise levels, we derive the analytical expressions of the GFP noise using the fluctuation-dissipation theorem (FDT) for the two models, which yield the decomposition of the total GFP noise (Method D). For the RC system, the square of the GFP total noise is given by

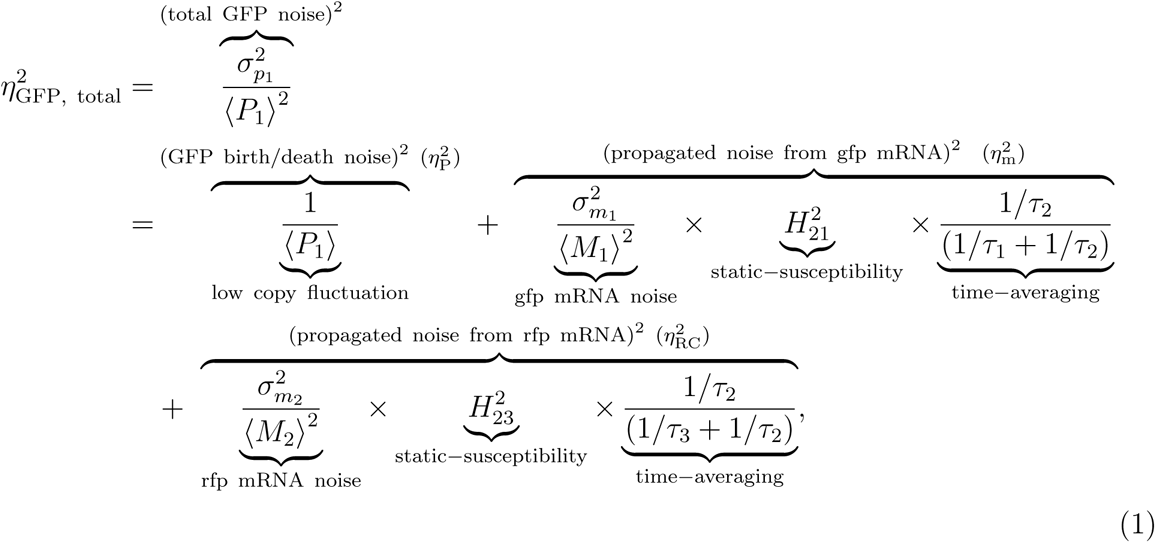

where the first two noise terms are the same as the ones in the UR case [Eq. (35) in Method D]. In particular, the first noise term represents the stochasticity from the random birth/death of protein (*η*_*p*_), which depends on the average number of GFP proteins. The second term is due to the fluctuations of the gene’s own mRNA (*η*_*m*_), which depends on the gfp mRNA noise and the contribution of gfp mRNA to its translation quantified by the susceptibility factor *H*_21_. The noise decomposition for the UR system with two noise terms is consistent with that revealed by previous studies [21, 30, 31]. The last term in Eq. (1) is specific to the RC system, which signifies the stochasticity from the other mRNA and depends on the rfp mRNA noise and the relative inhibition strength of gfp translation by rfp mRNA as characterized by the susceptibility factor *H*_23_.

To confirm that resource competition can lead to noise reduction, we compare the dependencies of the GFP noise on the parameter *J*_*p*_ that represents the translational capacity of limited resources in the host cell for the synthetic gene circuit, as shown by the solid light purple and blue dashed traces in Fig. 1d. It can be seen that GFP noise is always smaller in the RC case. As *J*_*p*_ increases its value, GFP noise in the RC case approaches the noise level in the UR case. This is reasonable because, as *J*_*p*_ approaches infinity, the terms in the RC model are reduced to those in the UR model, and the limitation on resources gradually disappears. It is worth noting that the changes in the GFP noise level with respect to *J*_*p*_ in the UR case are due to the parameter rescaling. We next find that the total GFP noise in the RC system first decreases then increases slightly with *J*_*p*_, as shown in Fig. 1h. This is due to the double-edged effects of resource competition on the noise: the noise reduction effect due to the resource limitation and generation of RC noise.

Figures 1h and 1d thus demonstrate how the noise composition changes with the resource availability. Specifically, the noise from the protein birth/death (*η*_*p*_) decreases with the resource availability (the yellow curve in Figs. 1h and 1d], due to the increased protein mean (the red curve in Fig. 1e). The noise due to the stochasticity from its own mRNA increases with the resource availability in the RC system (the maroon curve in Fig. 1h) due to the continuous relaxation of the resource limitations on translation, and approaches the level in the UR case, which does not change with translational capacity (the maroon curve in Fig. 1d). The RC noise decreases with the resource availability (the purple curve in Fig. 1h), as the inhibition of one mRNA to the translation of the other mRNA becomes smaller with more translational capacity. Taken together, our finding is that resource limitation can decrease gene expression noise but creates a new type of noise, creating a remarkable double-edged effect on gene expression noise.

### Elimination of resource competition noise through orthogonal resources

Orthogonal resources such as orthogonal RNAP and ribosomes have been developed to reduce unwanted couplings in gene circuits due to the competition for the host transcriptional and translational component [7–11]. We set out to study how the introduction of orthogonal resources affects the noise levels. With their addition into the two-reporter system, gfp and rfp now pull from two separate pools of RNAP and ribosomes, as shown in Fig. 2a. As a result, the inhibition links between the two genes caused by resource competition are removed while the resource constraint remains. We hypothesize that this would retain the noise decreasing effects of resource competition while nullifying the resource competitive noise.

**FIG. 2.**
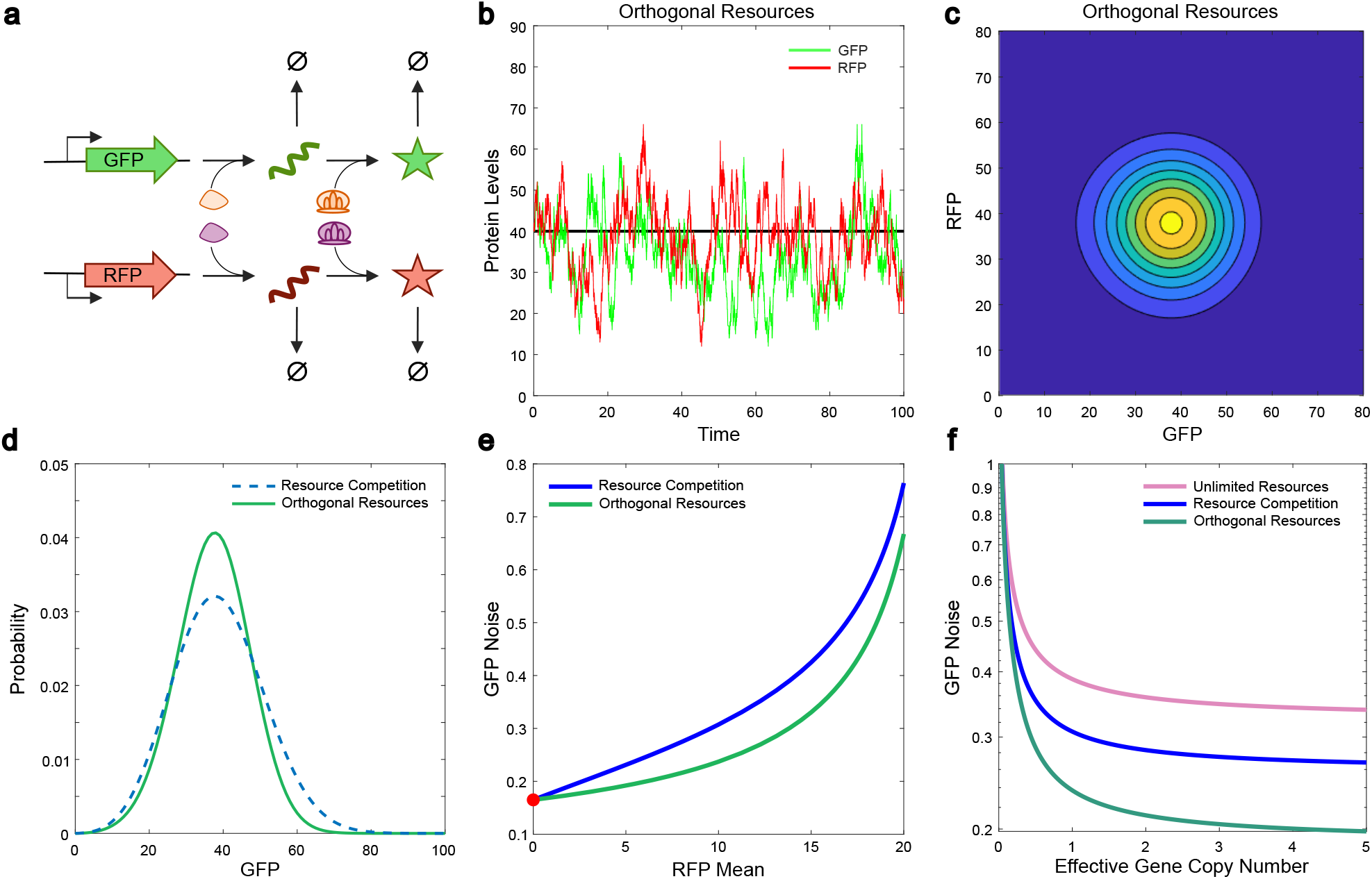
Elimination of resource competition noise through orthogonal resources. (a) Diagram illustrating how the use of orthogonal resources eliminates the competition for transcriptional and translational resources. (b) Gillespie stochastic trajectory of GFP (green curve) and RFP (red curve) expression in a two-gene circuit with orthogonal resources. (c) Distribution of GFP and RFP expression levels in a two-gene circuit with orthogonal resources obtained from the solutions of the master equation. (d) Distribution of GFP expression levels in a resource competition system (blue dashed curve) and an orthogonal resources system (green curve). (e) FDT analysis revealing the dependence of the GFP total noise levels on RFP mean. (f) FDT analysis giving the dependence of the protein total noise levels on the effective gene copy number in a system with unlimited resources (light purple curve), resource competition (blue curve), and orthogonal resources (green curve).

We construct a model for the orthogonal resource (OR) system [Eqs. (12) and (13) in Method A] compare its noise behavior with the RC system by rescaling the transcription and translation rate constants in the OR model to keep the same means of the mRNAs and proteins in the two models. The stochastic trajectories reveal that the expression levels of both proteins are no longer anticorrelated and fluctuate closer to the mean, as shown in Fig. 2b. Accordingly, the 2D GFP/RFP probability distribution is more compact near the mean, as shown in Fig. 2c. The probability distribution for the OR case (the solid green curve in Fig. 2d) is narrower than the RC case (the dashed blue curve in Fig. 2d), suggesting that utilization of orthogonal resources can reduce the protein noise levels. An analytical solution of the protein noise shows that the resource competitive noise has been removed from the system (Method D).

We then study how the GFP noise levels in the RC and OR systems change with the RFP mean by increasing the effective RFP gene copy number with the rescaled parameters in the OR system. As the RFP mean increases, more and more resources are taken by the RFP module, so the level of GFP protein mean linearly decreases, as shown in Fig. 2a, which is consistent with previous results [4]. With the rescaled parameters, we make the means of mRNAs and proteins in the OR system the same as the RC system. We find that, when there is no RFP, the noise levels in the two systems are equal, as shown by the red dot in Fig. 2e. As the RFP mean increases, the GFP noise levels in both systems increase because of the reduced copy number of GFP mRNA and protein (Fig. 2a), but the noise level in the OR system is always smaller (Fig. 2e) given that the RC system has the additional noise (*η*_*RC*_) and larger propagated noise from its own mRNA (*η*_*m*_), as shown in Figs. 2b and 2c. Note that the maximum RFP mean reached by increasing the copy number is due to the saturation mediated by resource limitation, as shown in Fig. 2d. This conclusion holds true if the plasmid copy numbers increase for both genes. The total noise decreases and reaches a saturation floor, as shown in Fig. 2f, which is consistent with the previous findings [32– 35]. Nonetheless, the noise levels in the UR system are always the largest, suggesting that utilizing orthogonal resources is a good strategy to eliminate the contribution of resource competition to gene expression noise.

### Control of gene expression noise by negatively competitive regulatory (NCR) controllers

Negative feedback has been utilized extensively to mitigate the effects of resource competition [12–18] and reduce gene expression noise [36–45]. We systematically study the noise attenuation ability of these negative feedback controllers in the context of resource competition. Previously, we proposed a controller topology for combating resource competition effects - negatively competitive regulatory (NCR) controller [18], as schematically illustrated in Fig. 3a. Briefly, in addition to their output proteins, each competing module also creates an sgRNA which is inhibitory towards the module that produced it. However, these sgRNAs cannot initiate inhibition until they complex with an inhibitory CRISPR moiety (e.g., dCas9), which are drawn from a fixed pool. The resulting inhibitory complexes can then initiate inhibition of their respective modules. This is an example of mRNA-mediated inhibition of transcription. Such a controller topology can be generalized to be placed at any one of four places in the protein production pathway, defined by whether the inhibition is mediated via mRNA or protein and whether the controller targets transcription or translation for inhibition. We henceforth define four controller subtypes, as shown in Fig. 3b: mRNA inhibitory transcription (MIX), protein inhibitory transcription (PIX), mRNA inhibitory translation (MIL), and protein inhibitory translation (PIL).

**FIG. 3.**
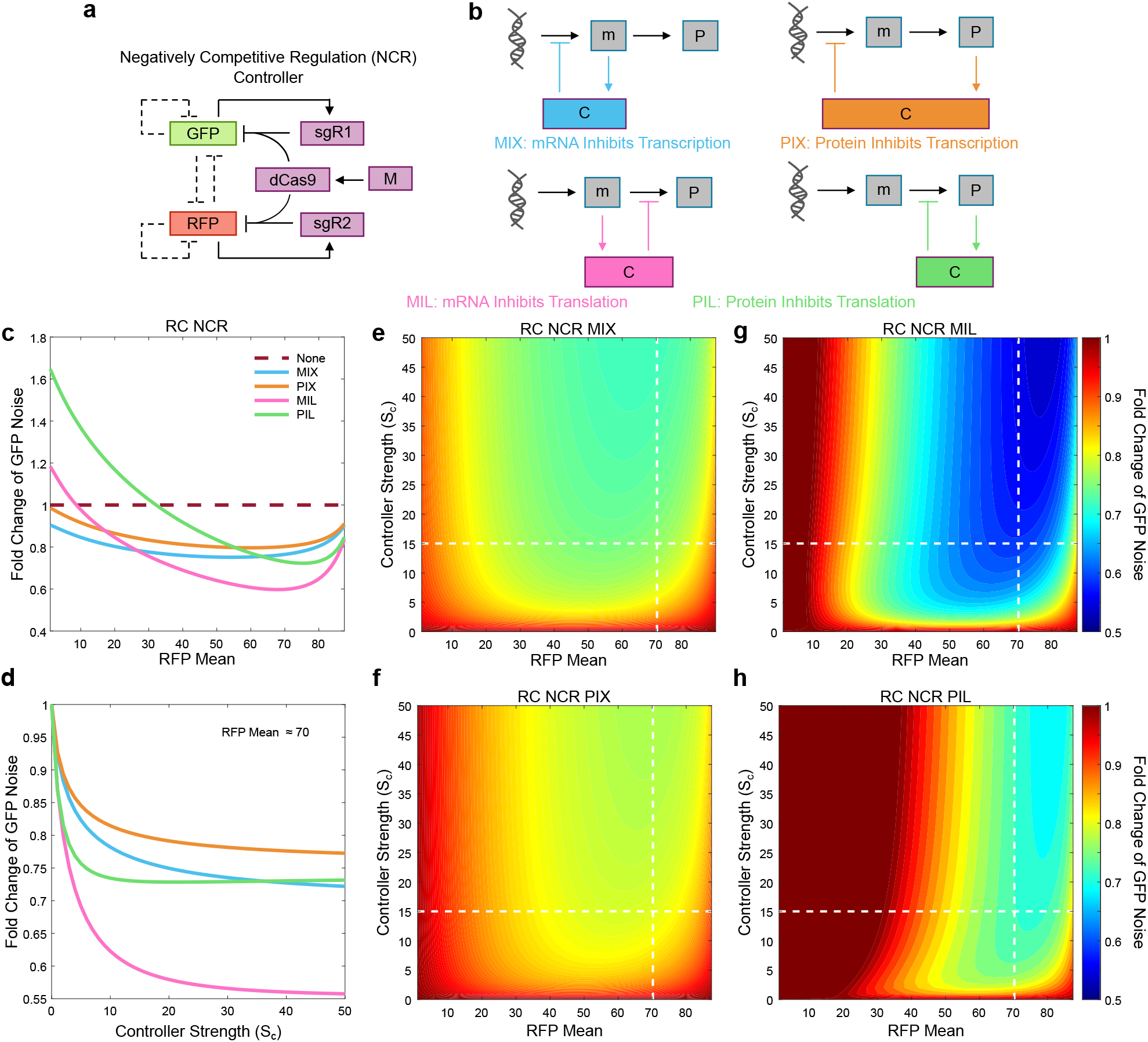
Control of gene expression noise by negatively competitive regulatory (NCR) controllers. (a) Topology of the NCR controller acting on a two-gene circuit. Each module (GFP, RFP) produces an inhibitory guide RNA (sgR1, sgR2) which is used for self-inhibiting complexes upon binding to a fixed pool of CRISPR moiety (dCas9). The blunt dash arrows show the self-inhibition and mutual inhibition between two modules due to resource competition. (b) General controller topologies applied at different positions in the protein production pathway defined by the inhibitory moiety type and the target of inhibition: mRNA inhibitory transcription (MIX), proteins inhibitory transcription (PIX), mRNA inhibitory translation (MIL), and proteins inhibitory translation (PIL). (c) FDT analysis demonstrating the normalized GFP total noise level on the RFP mean with NCR controllers applied in a resource competitive system for fixed controller strength *S*_*c*_ = 15. The noise levels are normalized to the base case without a controller. (d) The dependence of normalized GFP noise on *S*_*c*_ for RFP Mean fixed at 70. (e-h) Normalized GFP noise in the phase plane of RFP mean and NCR controller strength for NCR controller subtype (e) MIX, (f) PIX, (g) MIL, and (h) PIL. Deep blue color represents a strong decrease in the noise levels with respect to the base case with no controller, and deep red indicates the absence of a significant noise change or even a noise increase over the base case. The horizontal and the vertical white dashed lines represent the controller strength in panel (c) and the RFP mean used in panel (d), respectively.

To assess the noise reduction ability of these NCR controllers, we develop a generalized model for these controllers (Method E) and carry out FDT-based numerical analysis (Method F) for all the four subtypes with increasing RFP means for a fixed controller strength. Figure 3c shows the GFP noise normalized by the base case without any controller. For comparison, Fig. 3a shows the case of non-normalized GFP noise. We find that translation-inhibiting subtypes (MIL and PIL) have the largest noise reduction effects as RFP mean increases but they perform poorly at low RFP means where GFP noise even increases. Transcription-inhibiting subtypes (MIX and PIX) do not reduce noise significantly at high RFP means but are able to decrease noise consistently over all RFP mean values. Importantly, mRNA-mediated controllers (MIX and MIL) outperform those mediated by protein (PIX and PIL) in both the inhibition of transcription and translation cases. Figure 3d shows that normalized GFP noise decreases as the controller strength increases with a fixed large RFP mean but saturates beyond a certain point. Under a smaller or moderate RFP mean, normalized GFP noise in the PIL case can increase, as shown in Fig. 3b or first decreases then increases, as shown in Fig. 3c with increasing controller strength. These trends hold with other RFP means and controller strength. Figures 3e-3h demonstrate how GFP noise changes in the phase plane of RFP mean and controller strength for all controller subtypes, where deep blue regions represent large noise reduction whereas dark red regions indicate either little noise reduction or an increase in noise. MIL and PIL subtypes have both regions of deep blue but also dark red, indicating their increasing ability of noise reduction with RFP mean but poor starting performance at low RFP mean intervals. Heatmaps of MIX and PIX have neither much deep blue nor dark red colors, indicating their ability to consistently reduce noise over the entire RFP mean interval, albeit at moderate levels.

### Control of gene expression noise using local and global controllers

We investigate local and global negative feedback controllers that were used previously to mitigate resource competitive effects [15, 19]. Briefly, local controllers incorporate separate negative feedback loops, i.e., every module in the genetic system has its own negative feedback loop, as shown in Fig. 4a, but global controllers use a single shared negative feedback loop that represses all modules in the circuit, as shown in Fig. 4b. To compare the efficacy of these controllers in attenuating noise in the two-gene circuit, we perform FDT analysis for the systems with each controller applied utilizing one of the four placement subtypes. We find that global controllers perform poorly. First, the Global MIX and PIX systems reduce the noise slightly at low RFP means but are barely able to reduce noise at higher RFP means, as shown in Figs. 4c and 4a. Second, the global MIL and PIL systems significantly increase noise for most of the RFP mean intervals, as shown in Figs. 4d and 4b. This finding is consistent with the global controller’s inability to attenuate the winner-takes-all resource competitive behavior [18].

**FIG. 4.**
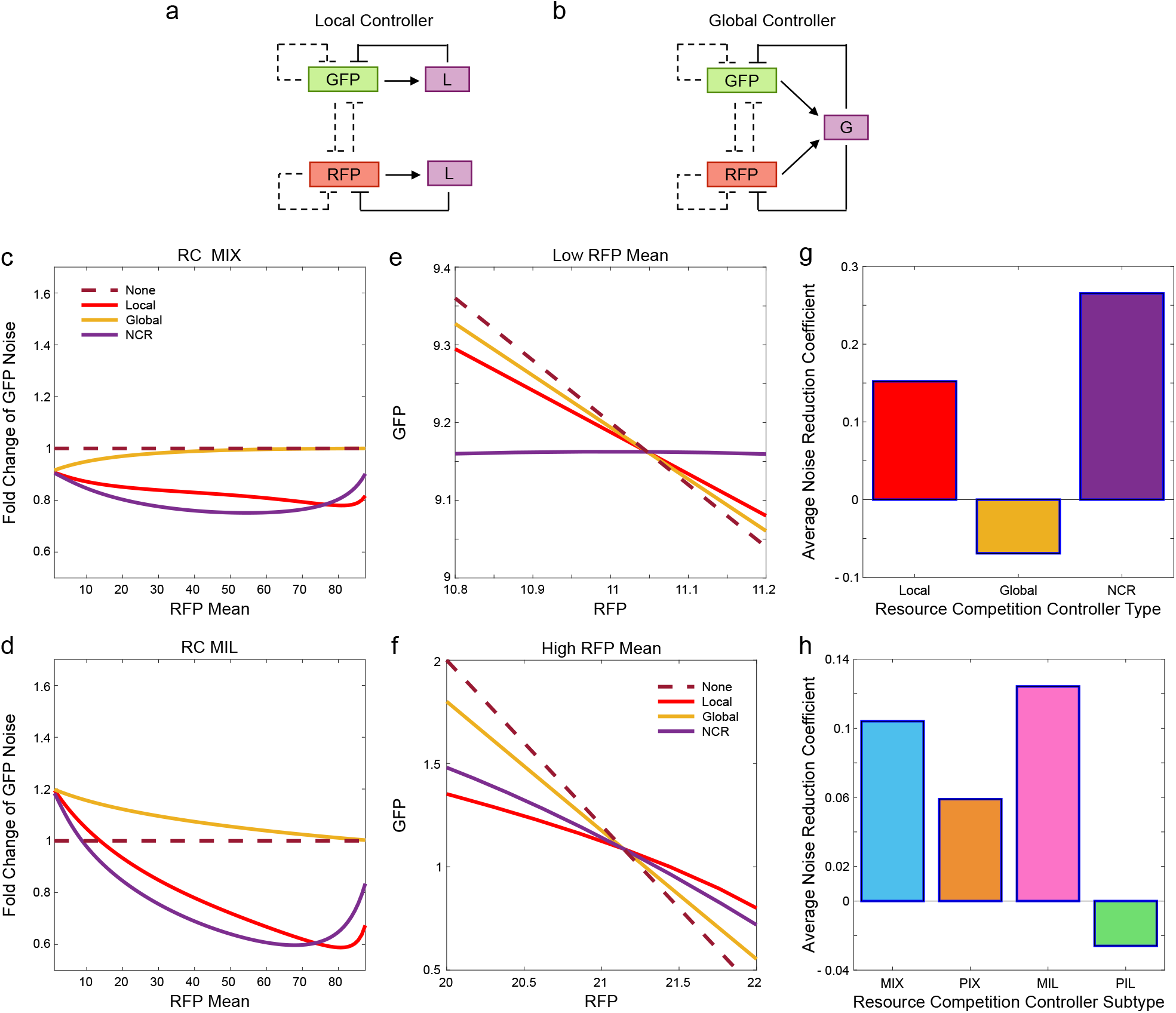
Control of gene expression noise using local and global controllers. (a, b) Diagrams of the local and global controller, respectively, acting on the two-gene system. (c, d) Dependence of the normalized GFP noise levels with local controllers (red curves), global controllers (yellow curves), and NCR controller (purple curves) using, respectively, MIX and MIL subtypes. The noise levels are normalized to the base case with no controller. (e, f) Deterministic dependence of GFP level on RFP level with various MIL controllers applied, where the parameters are fixed to intersect just at one point for low and high RFP mean, respectively. (g) Average noise reduction coefficient for local, global, and NCR controller types. (h) The coefficient for controller subtypes MIX, PIX, MIL, and PIL (h).

Systems with a local controller perform more poorly than NCR controllers at low/moderate RFP means, as shown in Figs. 4c, 4d, 4a, and 4b. However, there exists a critical RFP mean value for which Local and NCR cross in efficacy. The value of this cross-point is parameter-dependent, but NCR typically performs better at low/moderate RFP means while a local controller typically performs better at high RFP means. The reason for this cross in controlling efficacy can be seen from the GFP versus RFP mean graphs, as shown in Figs. 4e and 4f. As the RC noise in GFP is qualitatively related to the slope at a given point on the graph, a shallower slope on a GFP versus RFP mean graph indicates weaker noise. At low/moderate RFP means (Fig. 4e), the two-gene system with the NCR controller applied has a rather flat slope, indicating that the NCR controller can nearly decouple the two genes from their resource competition at this point and thus is better at noise reduction. However, at a high RFP mean (Fig. 4f), the slopes of the curves with NCR and local controller interchange, with the local controller curve now having the shallowest slope and thus a better noise reduction capability.

To obtain more general and conclusive results regarding which controller and placement subtype are optimal for noise reduction, we define an average noise reduction coefficient as the average fold change in noise across all RFP means, controller strengths, and/or controller types/subtypes (Method G). The average noise reduction coefficients are shown in Fig. 4g for each controller type and in Fig. 4h for all three controller types. These results demonstrate that the global controllers and the PIL controllers on average perform poorly (given their negative noise reduction coefficients) and are generally inappropriate for building an efficacious noise reducing system. Further support for this finding is presented in Figs. 4g-4j, where a deep red color emerges for the majority of the phase plane of RFP mean and control strength for the global controller, and in Figs. 4f and 4j for the PIL subtypes. While Local and NCR systems are both relatively efficacious in attenuating noise, NCR is significantly more consistent in its ability to reduce noise, as shown in Fig. 4g. The two best performing placement topologies are the MIX and MIL subtypes, with the MIL placement more consistent in its ability to reduce noise (Fig. 4h). Furthermore, it can be concluded that a system’s noise reduction capability in general is determined more by the type of controller than by which controller subtype chosen. This can be seen as the noise reduction coefficients from altercation of controller type (Fig. 4g) span approximately 0.3 units, whereas altercating the controller subtype only results in a smaller span of approximately 0.14 (Fig. 4h).

### Control of gene expression noise through combined negative feedback controllers and orthogonal resources

Having demonstrated that both orthogonal resources and negative feedback controllers can attenuate noise, we investigate whether their combinations can improve the noise-control capability. We first focus on the combinations of the most effective controllers in the RC system, including local-MIX, local-MIL, NCR-MIX, and NCR-MIL, with orthogonal resources. We then determine whether the application of an OR system can benefit negative feedback controllers for noise reduction. As shown in Figs. 5a and 5a-5c, the GFP noise normalized to the RC base case without a controller indicates that the use of orthogonal resources consistently benefits local-MIX and NCR-MIX controllers. However, using OR system is barely beneficial to local-MIL and is deleterious to NCR-MIL, as shown in Fig. 5b. The GFP noise normalized to the OR case without a controller reveals that the addition of a local-MIX or NCR-MIX controller is consistently beneficial for the OR system in further attenuating noise, as shown in Figs. 5c and 5d-5f. Even though local-MIL and NCR-MIL controllers do not provide any benefit for the OR system at small RFP mean in reducing noise, the synergy emerges in large RFP mean intervals. There is thus consistent synergy between the local/NCR controllers and orthogonal resources.

**FIG. 5.**
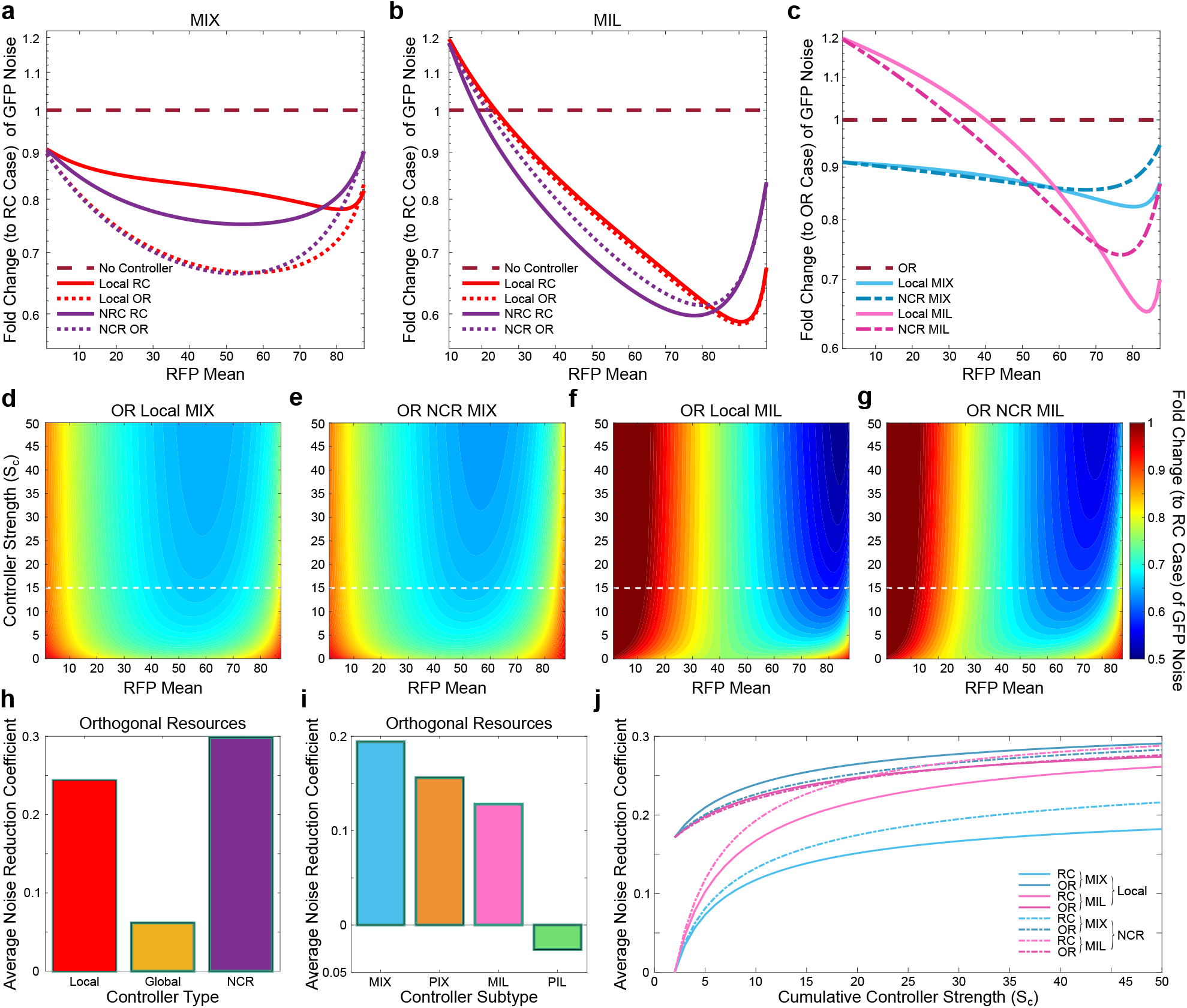
Control of gene expression noise through combined negative feedback controllers and orthogonal resources. (a, b) Dependence of the normalized GFP noise levels on RFP mean with a local controller (red curves) and NCR controller (purple curves) combined with orthogonal resources (solid curves), respectively, for MIX and MIL controller subtypes. The dashed curves are cases without orthogonal resources. All data are normalized to those of the system without a controller (maroon dashed curve). (c) Dependence of the normalized GFP noise levels on RFP mean for the OR system combined with a local controller (solid curves) or an NCR controller (dashed curves) for either the MIX (blue curves) or MIL (fuchsia curves) controller subtypes. All data are normalized to those in the case with orthogonal resources but no controller (maroon dashed curve). (d-g) Normalized GFP noise in the phase plane of RFP mean and controller strength, respectively, for local-MIX, NCR-MIX, local-MIL, and NCR-MIL combined with orthogonal resources. Deep blue color represents regions of higher noise reduction. The horizontal white dashed lines represent the controller strength shown in panels (a) and (b). The average noise reduction coefficient for (h) different local, global, and NCR controllers with orthogonal resources, and (i) different controller subtypes: MIX, PIX, MIL, and PIL with orthogonal resources. (j) Dependence of the cumulative average noise reduction coefficient on the controller strength for the cases with MIX and MIL of NCR/local controllers applied with or without orthogonal resources.

Heatmaps of each of the four controller subtypes over the RFP mean controller strength plane demonstrate similar behaviors for a range of controller strength, with the fold changes in GFP noise becoming more pronounced as controller strength increases, as shown in Figs. 5d-5g. Both combinations of an OR system with Local-MIX and NCR-MIX are the most effective at reducing noise when compared to a RC system without control. However, the OR local-MIL and OR NCR-MIL combinations possess a remarkable ability to reduce noise for higher RFP mean values.

To find a general synergy between the negative feedback controllers and orthogonal resources, we calculate the average noise reduction coefficients for all controller types and subtypes in the OR system, as shown in Figs. 5h and 5i. Similar to the system without orthogonal resources, the global controller and the PIL subtype perform poorly, though the global controllers (MIX and PIX subtypes) are a much better contender when added to the OR system (Figs. 5h, 4a, 4b, 5b, and 5e). Furthermore, the local and NCR controllers perform well, with NCR being on average more consistent in its noise-reduction capability. However, the order of the most efficacious controller subtype is different when applied to an OR system than the RC system, as shown in Figs. 5i and 4h, where transcriptional inhibition subtypes (MIX and PIX) demonstrate a better ability to reduce noise than translational inhibition subtypes (MIL and PIL). Figure 5j shows the dependence of the accumulative noise reduction coefficient on the controller strength for the eight best controller and subtype combinations. The curves in the OR systems have a much higher starting point at small controller strength range due to the strength-independent noise reduction by orthogonal resources. The NCR controllers are always better than local controllers in the RC systems, while in the OR system, the latter are slightly better than the former. This is reasonable as the NCR controller is specifically designed to mitigate the effects of resource competition. The addition of orthogonal resources significantly improves the less efficacious NCR-MIX and local-MIX controllers but does not enhance the maximum noise-reduction capability for the local-MIL and NCR-MIL controllers. It is worth noting that, although the noise reduction coefficient of the PIX subtypes is greater than that of MIL subtypes here (Fig. 5i), this effect is largely due to the fact that the global-PIX controller is significantly improved (Fig. 5b). That is the reason we choose to analyze MIL over PIX in the best cases comparison in Fig. 5j despite PIX’s seemingly higher noise reduction coefficient.

## DISCUSSION

To uncover and understand the origin of gene expression noise is a fundamental problem in systems and synthetic biology and has been investigated extensively in the past [21–28]. However, in all the existing models, unlimited cellular resources are assumed, which is unrealistic for synthetic gene circuits with multiple coactivated modules. Our present work has demonstrated quantitatively that resource competition can significantly affect the noise behavior of synthetic gene circuits. A key finding is that resource competition has a double-edged effect on protein expression noise levels. In particular, resource competition is able to reduce noise by applying resource constraints to the system, in the absence of which the gene expression noise level is maximized. The noise-reduction capability can be attributed to self-inhibition introduced indirectly into the system by resource competition. However, the competition introduces a new type of noise (resource competitive noise) as it creates anticorrelated links between the gene modules. The double-edged effect makes the dependence of the total noise level on the resource availability strikingly non-monotonic. Incorporation of orthogonal resources can take advantage of this effect to reduce the noise by orthogonalizing the resource constraints on each gene module. This technique keeps the resource constraints but removes the effects of inter-module resource competition, allowing for fewer variables affecting gene expression while still using self-inhibition to keep the expression levels relatively close to the mean. Development of a completely insulated OR system in the future has the potential to significantly improve the control of resource competition.

From a control perspective, negative feedback loops are often used to suppress the noise level in gene expression [36–45]. However, previously none of the methods took into account resource competition. Nonetheless, a number of negative feedback controllers such as local, global, and NCR controllers have been used in the past to negate unwanted effects of resource competition [12–18], raising the question of whether these controllers can be used to reduce gene expression noise in the circuits with limited resources. Our analysis reveals that global controllers are typically quite ineffective at reducing noise but often can increase it. In a recent work, we demonstrated that global controllers are not efficacious at reducing the effects of winner-takes-all type of resource competition [18], implying its inability to reduce resource competitive noise. Local and NCR controllers, however, both are efficacious at reducing system noise, with the latter outperforming the former at low RFP means and the opposite behavior at high RFP means. A useful result is that the controller placement topology within the protein biosynthesis pathway can drastically alter the noise reduction ability of a controller. Particularly, we find that inhibition mediated via mRNA (MIX and MIL subtypes) is more efficacious than inhibition mediated by protein (PIX and PIL subtypes). Combining orthogonal resources systems and negative feedback controllers makes the transcriptional inhibition strategies (MIX and PIX subtypes) more effective than translational inhibition strategies (MIL and PIL subtypes).

It is important to note that the large array of negative feedback controller topologies analyzed in this work have biological underpinnings and are not merely theoretical. For example, the MIX, MIL, and PIX subtypes of the local controller have been demonstrated in vivo by various groups [15, 17, 46], the local-PIL controller can be constructed via expression of orthogonal, sequence-specific RNA-binding proteins as inhibitory effectors [46, 47], and a global controller architecture can be constructed from each of these systems by replacing the orthogonal feedback modules with copies of the same regulator [19] where each module performs the same negative feedback operation but these production pools are shared amongst modules. While the NCR controller type is the newest proposed negative feedback architecture, it has not yet been synthetically constructed in vivo models. In our recent work [18], we demonstrated how an NCR-MIX controller could be theoretically constructed using inhibitory deactivate dCas systems. It is important that careful consideration should be given to which dCas system is incorporated in such a design as many dCas systems suffer from the problem of strong/irreversible dCas moiety binding [48]. This NCR-MIX controller assumes a constant production of Cas moiety while individual genetic modules produce sgRNAs that mediate repression. Numerous dCas systems have been reported to demonstrate sequence-specific RNA-binding capabilities, opening up the potential for in vivo MIL constructs for the NCR controller type [49]. Many of these systems also exhibit sequence-specific DNA-binding behavior. However, recently a few systems have been discovered that naturally target RNA such as Csm3, Cmr4, Csm6, and Csx1, the extensive family of Cas13 proteins [50, 51]. In addition, complex signal processing functions have been designed utilizing adaptive zinc finger protein complexes (e.g., Bashor et al. demonstrated that a generalized system utilizing sequence-specific zinc-finger proteins (ZFPs) and a clamp composed of PDZ moieties strung together which accepts these ZFPs is capable of performing very complex signal processing [52]). Since complexing between a conserved moiety (PDZ clamp) and sequence-specific moieties (ZFPs) is required to form the inhibitory complexes, it is possible to generate NCR-PIX systems. Furthermore, certain ZFP constructs have been demonstrated to have sequence-specific RNA-binding capabilities: such RNA-binding ZFPs can be potentially paired with PDZ/PDZ-like clamps to open up the possibility of NCR-PIL systems [53].

In our work, potential sources of extrinsic noise in the system such as the fluctuation of the copy numbers of the transcriptional resource (RNAPs) [35] and translation resource (ribosomes) are not included. It was reported that sharing a common regulator pool could result in indeterminacy of extrinsic noise [54]. It is worth noting that resource competition makes the two-reporter expressions anticorrelated. Significant extrinsic noise making two reporters fluctuate in a positively correlated fashion, together with strong resource competition, may lead to a circular 2D probability distribution. Our models do not take into account the bursting of gene expression, which can be simulated with a two-state model [26, 55] and can lead to a significant noise at the transcriptional level, especially under the context of a limited level of RNAPs. The contribution of these factors to the noise of synthetic gene circuits needs to be characterized. Furthermore, our noise analysis has been carried out using the most basic resource competitive system: an unregulated, unlinked two-gene system. For future investigation, we intend to look at the noise behavior in more complex and dynamic systems, such as the dual self-activating or cascading circuits that we have analyzed recently [6, 18]. The noise from other circuit-host interactions such as growth feedback [56– 61] adds another layer of complexity to the stochastic gene expression of synthetic circuits. The stochasticity in cellular growth and its propagation to the synthetic gene circuits is another potential source of fluctuations [61–64]. It may also prove insightful to investigate the noise-reducing effects of other methods for attenuating resource competitive effects, such as incorporating incoherent feedforward loop topologies [12, 16, 65] or antithetic integral feedback [66, 67] into the circuit.

## ACKNOWLEDGEMENTS

This project was supported by NIH grant (R35GM142896 to X.-J.T.). H.G. and A.S. were also supported by the Arizona State University Dean’s Fellowship. Y.-C.L. was supported by Office of Naval Research under Grant No. N00014-21-1-2323.

## METHODS

### A. Models for systems without controllers

Two identically regulated genes are placed within the same circuit, as shown in Fig. 1(a). The first gene results are shown through GFP protein expression, while the second uses RFP protein. Three separate scenarios are considered for this gene circuit: unlimited resources (UR), resource competition (RC), and orthogonal resources (OR). The system is generally described by a set of ordinary differential equations:

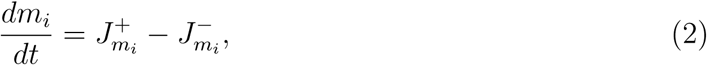

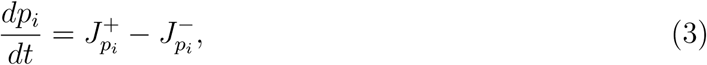

where *m*_*i*_ and *p*_*i*_ represent the mRNA and the protein concentrations, respectively, with the subscripts *i* = 1, 2 indicating the first or the second mRNA/protein (GFP, RFP), 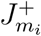 and 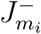 are respectively the production and degradation rates of mRNAs, and the respective rates for proteins are 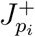 and 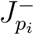. The degradation rates of mRNAs and proteins are proportional to their concentration: 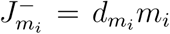 and 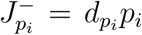, where 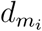 and 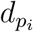 are constants of degradation. The forms of 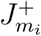 and 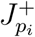 depend on the assumptions used in each particular system.

A system with unlimited resources (UR) has absolutely no competition and can be modeled by

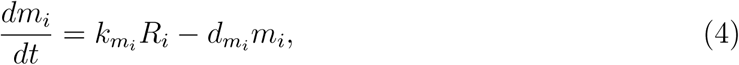

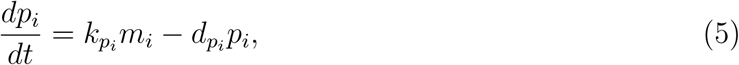

where 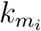 and 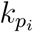 are the production rate constants, 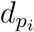 and 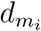 are the degradation rate constants for the mRNAs and proteins, respectively, and *R*_*i*_ represents the effective plasmid DNA copy number. We thus have 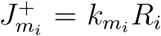 and 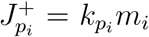. Setting these derivatives to zero gives the steady-state mRNA and protein mean concentrations, respectively, as

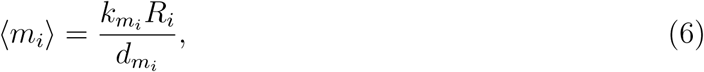

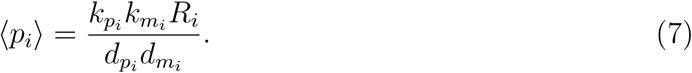

This type of model has been widely used to simulate gene expression levels and the corresponding noise [21–28].

A system with limited shared resources causing resource competition (RC) stipulates the genes compete with both themselves and each other for the limited transcriptional and translation resources. The equations for this system [6] are

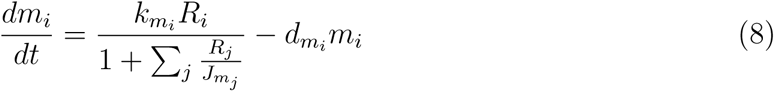

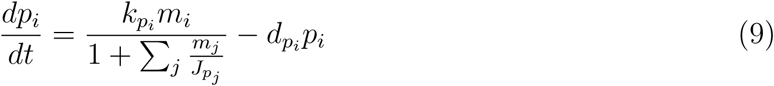

where 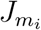 and 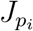 are the effective transcriptional and translational capacities of limited resources in the host cell for synthetic gene circuits, respectively. Note that as 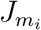 and 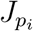 approaches infinity, the RC model reduces the UR model, so

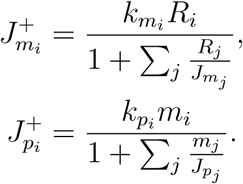

Again, setting these derivatives to zero yields the steady state mRNA and protein mean concentrations as

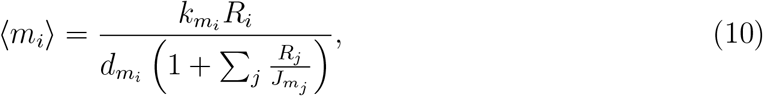

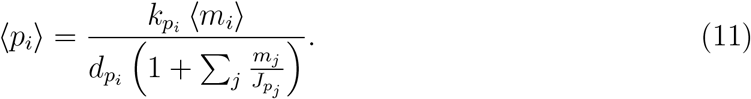

A system with orthogonal resources (OR) eliminates the resource competition between genes, where the two genes still compete with themselves due to the limitation in the resources but not with each other. The system is described by

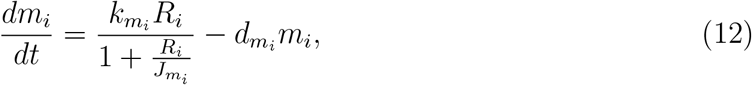

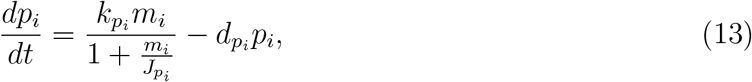

with 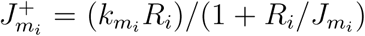 and 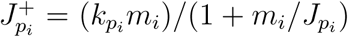. In the steady state, the mRNA and protein mean concentrations are, respectively,

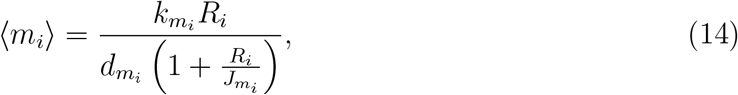

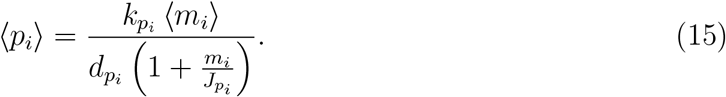

### B. Gillespie stochastic simulations

To simulate the stochastic trajectories of mRNA and protein levels, we use the standard Gillespie stochastic simulation method [29], with the following steps.

1. Define the system according to expression levels using the following matrices: 𝒮 - the number of each molecular used as reactants and 𝒫 - the number of each molecular made as products :

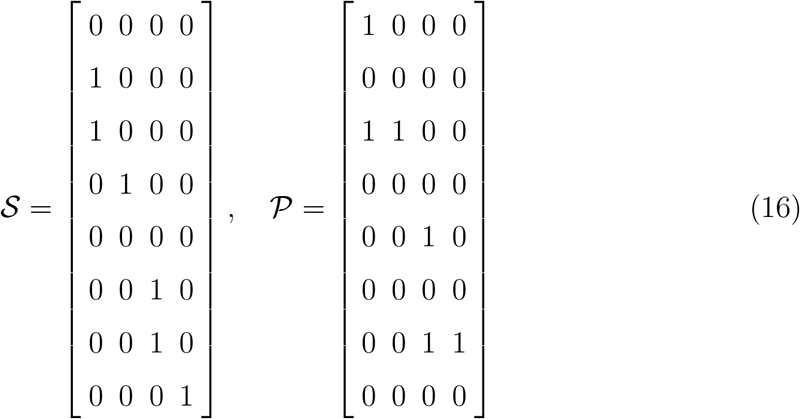
2. Initialize **n** as the vector of the mean copy numbers of mRNAs and proteins, **n** = [*M*_1_, *P*_1_, *M*_2_, *P*_2_] and set time *t* = 0. It is noted that we use capitalized letters *M*_*i*_, *P*_*i*_ to denote the numbers of mRNAs and proteins, which can be converted to the concentrations *m*_*i*_ = *M*_*i*_*/*Ω and *p*_*i*_ = *P*_*i*_*/*Ω by using a system size factor Ω.
3. Update the reaction rate vector **v** using current **n**:

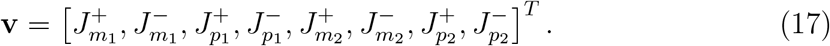
4. Calculate the reaction probability vector:

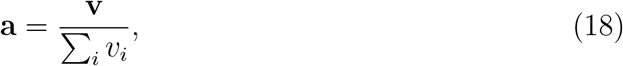

where *v*_*i*_ is the *i*th component of **v**.
5. Reaction *r* occurs if

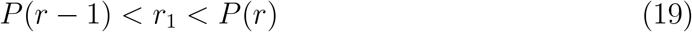

and the time step to the next reaction is

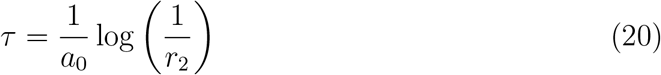

where *a*_0_ = ∑_*i*_ *v*_*i*_, 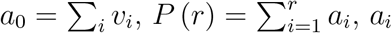 is the *i*th component of **a**, *r*_1_ and *r*_2_ are two uniformly distributed random numbers from the unit interval [0, 1].
6. Update **n** and *t* according to

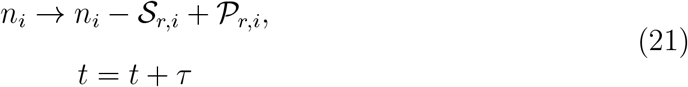
7. Continue steps 3-6 until *t* reaches a predetermined maximum time.

### C. Master equation

The probability *P* (*M*_1_, *P*_1_, *M*_2_, *P*_2_, *t*) of the system in the state (*M*_1_, *P*_1_, *M*_2_, *P*_2_, *t*) at time *t* is governed by the master equation:

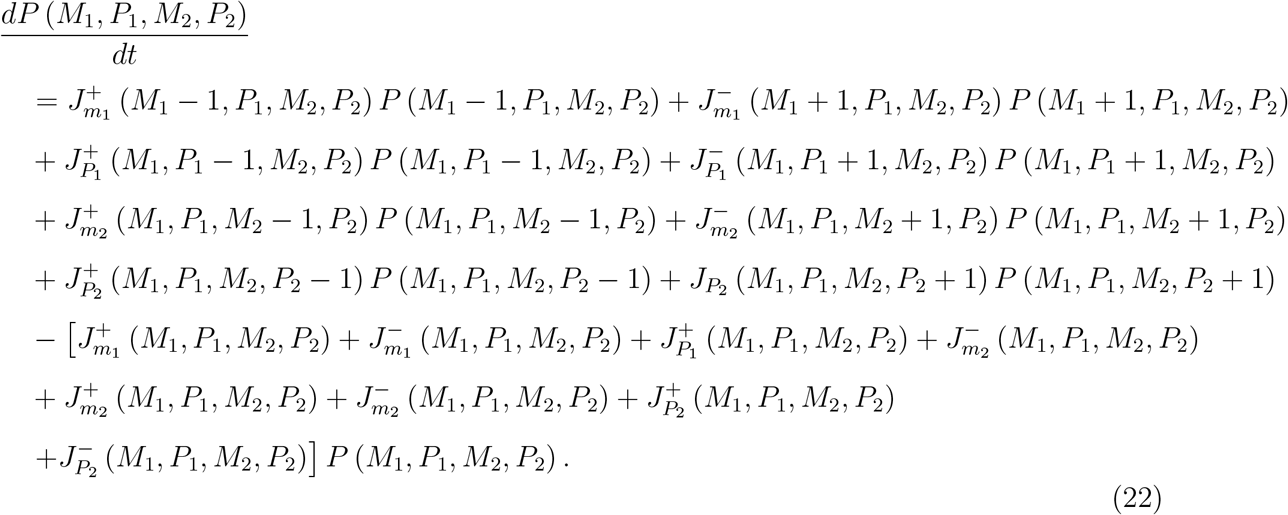

Equation (22) can be written in matrix form as

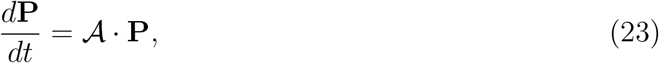

where **P** is the state probability vector, and 𝒜 is the transition rate matrix from state (*M*_1_ + *i, P*_1_ + *j, M*_2_ + *k, P*_2_ + *l*) to state (*M*_1_, *P*_1_, *M*_2_, *P*_2_), which is defined as

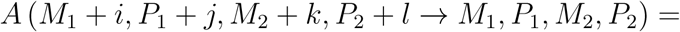

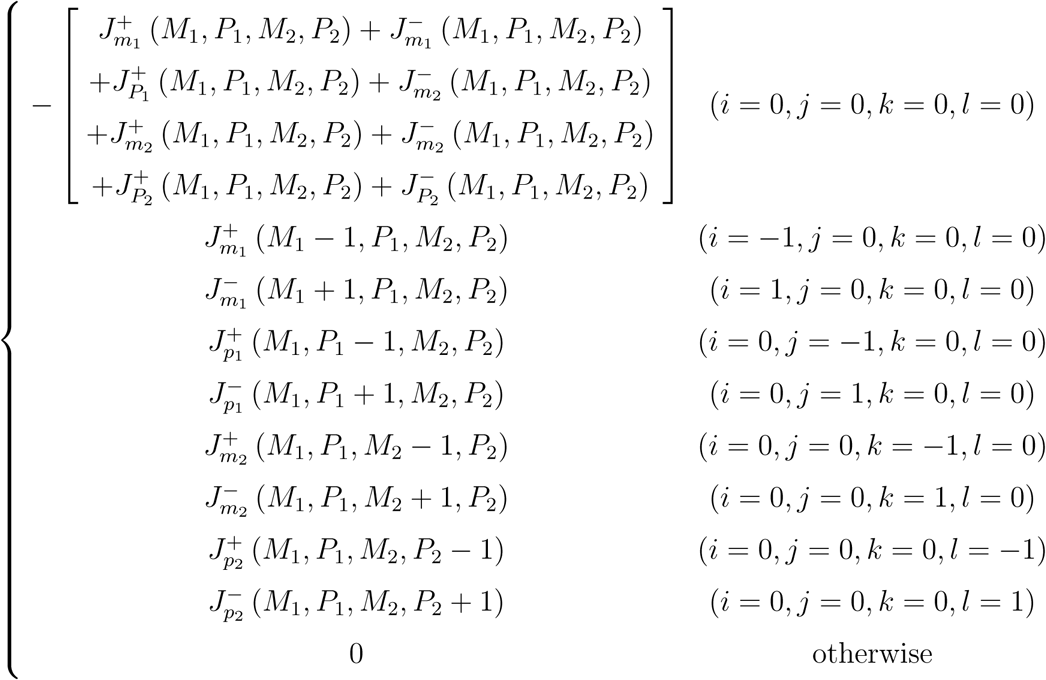

We solve the master equation until the system reaches a steady-state distribution. The boundary is set to [0, 3 ∗ *µ*_*i*_] as that the probabilities at right and top boundaries are small. *µ*_*i*_ = {⟨ *M*_1_⟩, ⟨*M*_2_⟩, ⟨*P*_1_⟩, ⟨ *P*_2_⟩} is the mean number of mRNAs or proteins. The no-flux boundary conditions are used to conserve probability. The master equation is used in Figs. 1d, 1g, 2c, and 1b.

### D. Analytical solution from the normalized fluctuation-dissipation theorem

We define the noise (*η*) for both mRNAs and proteins as the coefficient of variation:

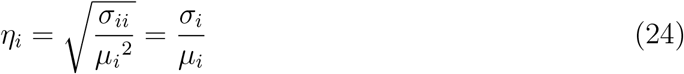

where 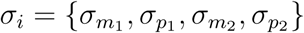 is the standard deviation (*σ*_*ii*_ as variance).

To obtain the analytical expressions of gene expression noise, we use the normalized version of the FDT [31]:

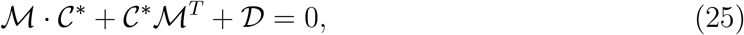

where 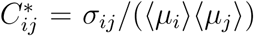 is the normalized covariance, *M*_*ij*_ = − *H*_*ij*_*/τ*_*i*_ is the elements of the dynamical matrix with

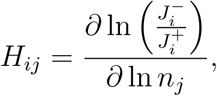

and *D*_*ii*_ = 2*/*(*n*_*i*_*τ*_*i*_). Here, *n*_*i*_ with *i* = {1, 2, 3, 4} represents the copy number of *m*_1_, *p*_1_, *m*_2_ and *p*_2_, respectively.

For the resource competition (RC) system, the matrices ℳ and 𝒟 are defined as

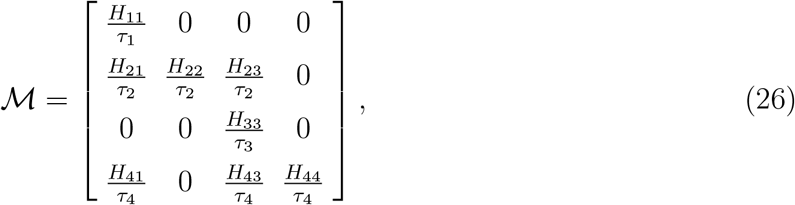

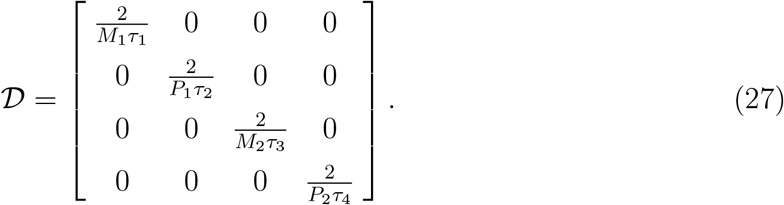

The normalized FDT equation can be solved to yield the following noise levels of mRNAs and proteins:

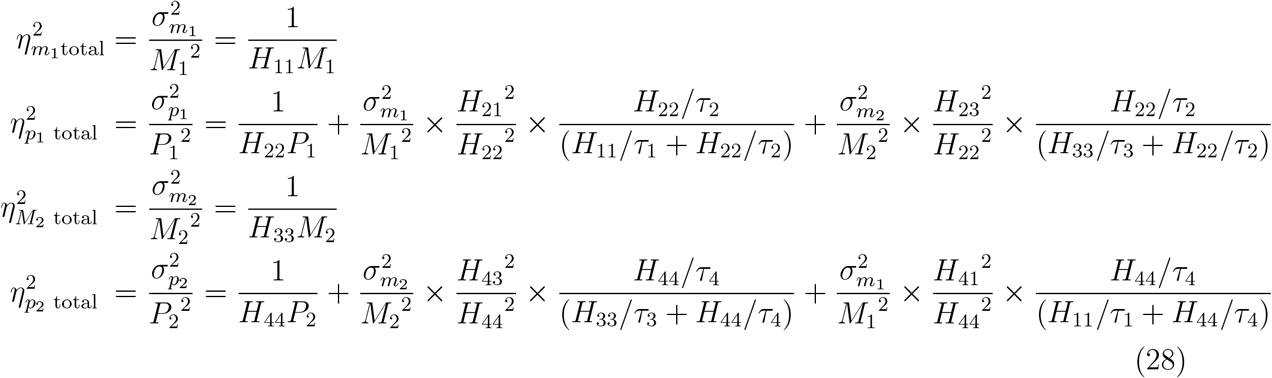

with

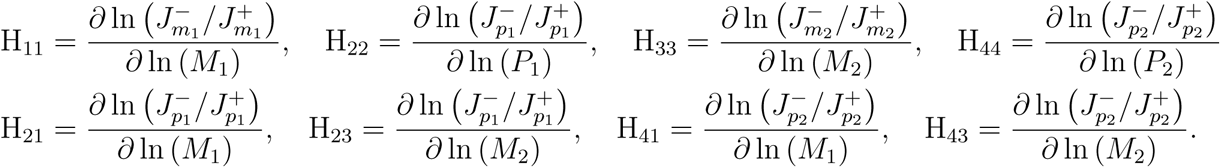

The revised production and degradation rates taking into account the system size factor Ω are

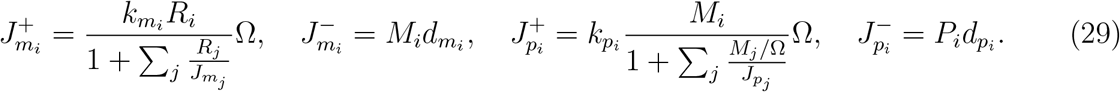

Substituting the parameter values of our model *H*_11_ = 1, *H*_22_ = 1, *H*_33_ = 1 and *H*_44_ = 1 into Eq. (28), we get

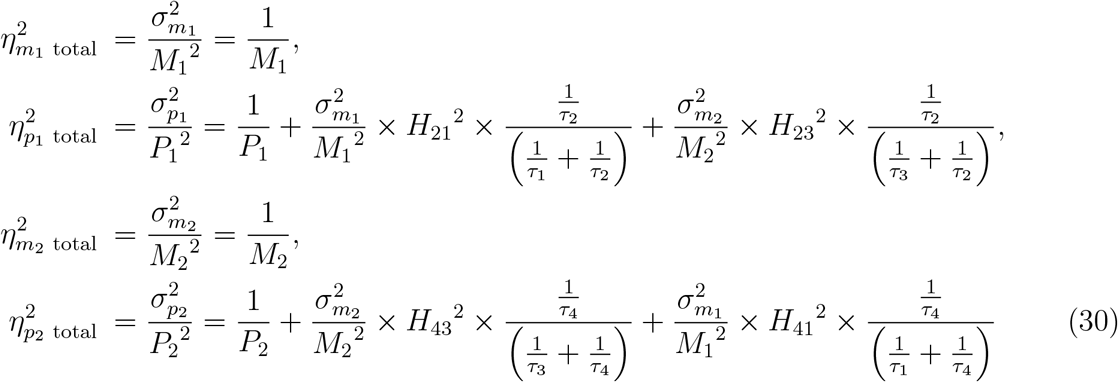

For the unlimited resources (UR) and the orthogonal resources (OR) systems, the matrices ℳ and 𝒟 are given by

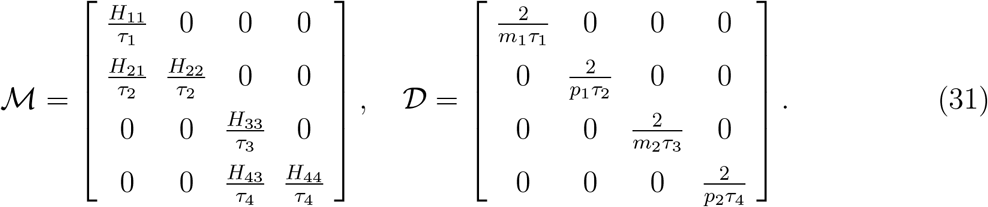

We then solve the normalized FDT equations to obtain

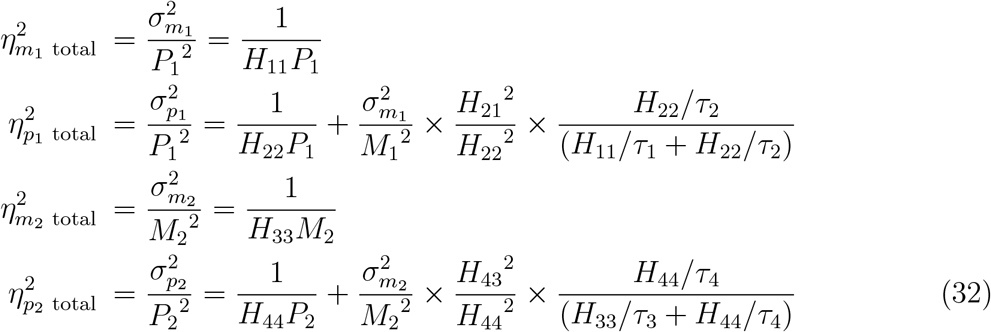

with

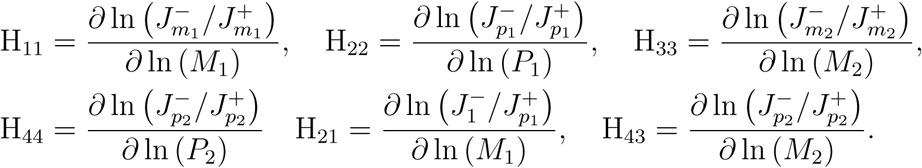

For the unlimited resource (UR) system, we have

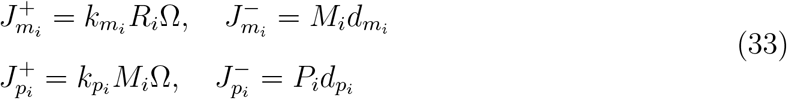

For the orthogonal resource (OR) system, we have

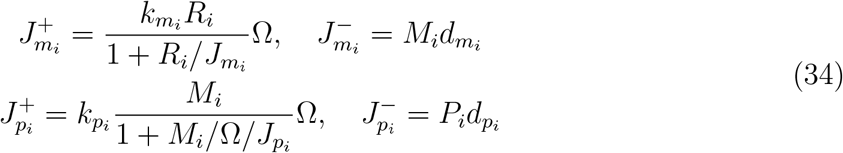

Substituting *H*_11_ = 1, *H*_22_=1, *H*_33_ = 1, and *H*_44_ = 1 into Eq. (32), we obtain

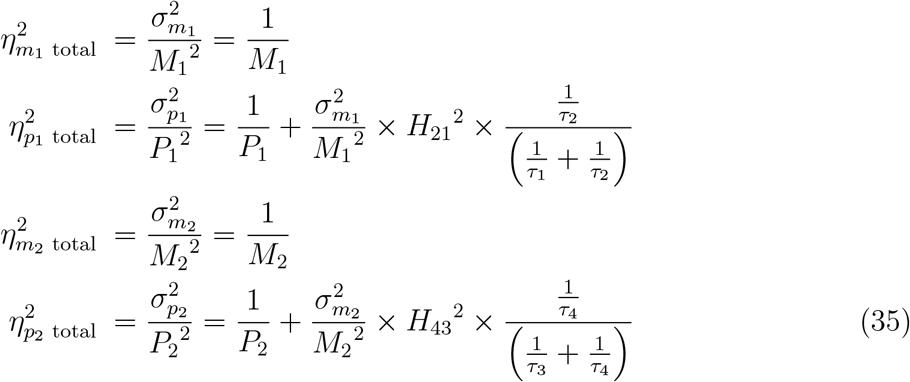

### E. Generalized model with negative feedback controllers

We consider three types of negative feedback controllers to mitigate the resource competition effects: local, global, and negatively competitive regulator (NCR) controllers. For each type of controller, depending on the mediators and targets of the negative feedback, there are four subtypes: mRNA inhibiting transcription (MIX), protein inhibiting transcription (PIX), mRNA inhibiting translation (MIL), and protein inhibiting translation (PIL). To model the noise resulting from embedding a negative feedback controller into the two-gene circuit, we modify the previous model [18] by including the translational step in protein biosynthesis. The system with both resource competition and control is described by

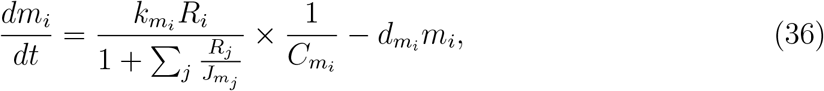

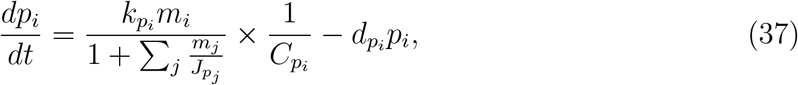

where 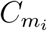 and 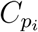 characterize the repression of transcription and translation by the controller, respectively, which are given by

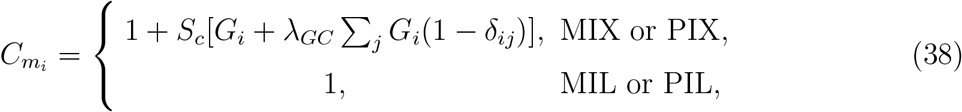

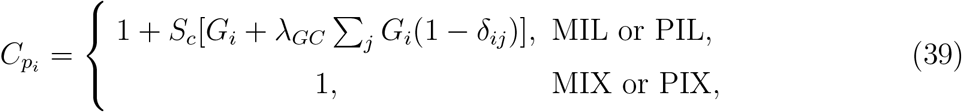

where *S*_*c*_ is the strength of the controller, *G*_*i*_ represents each gene’s contribution to the controller’s negative feedback activity, which is given by

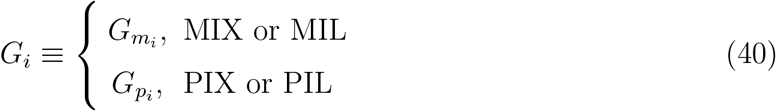

with 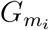 and 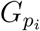 defined as

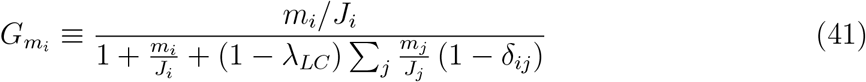

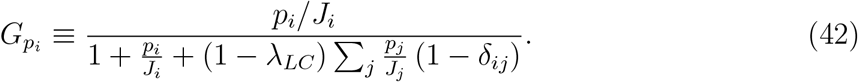

In Eqs. (41) and (42), *δ*_*ij*_ is the Kronecker delta matrix, where *δ* = 1 for *i* = *j* and zero otherwise, and the binding affinity of the repressive complex is represented by *J*_*i*_. The hyperparameters *λ*_*LC*_ and *λ*_*GC*_ in Eqs. (38), (39), (41), and (42) are given by

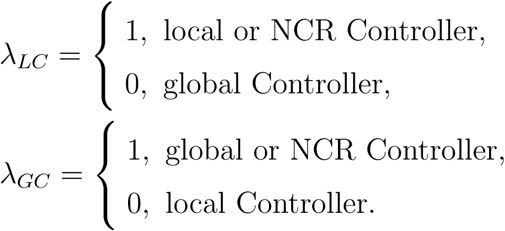

Similarly, the systems with orthogonal resource and controllers are described by

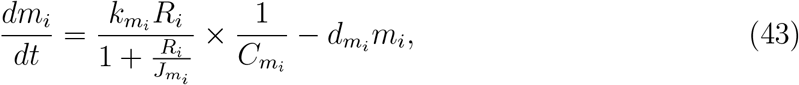

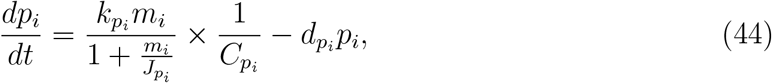

where 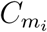 and 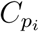 are given by Eqs. (38) and (39), respectively. Equations (36-44) model all twenty-four negative feedback and mixed controller types studied.

### F. Determining gene expression noise levels using fluctuation-dissipation theorem

To calculate the mRNA and protein noise levels with negative controllers, we use the fluctuation-dissipation theorem [30, 31]:

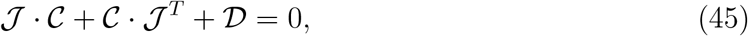

where 𝒞 is the correlation matrix with the covariance *σ*_*ij*_ and 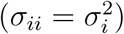, 𝒥 is the Jacobian matrix of all partial derivatives of the vector field, and 𝒟 is the diffusion matrix. With **n** as the vector of molecule copy numbers and 𝒩 as the stoichiometric matrix given by

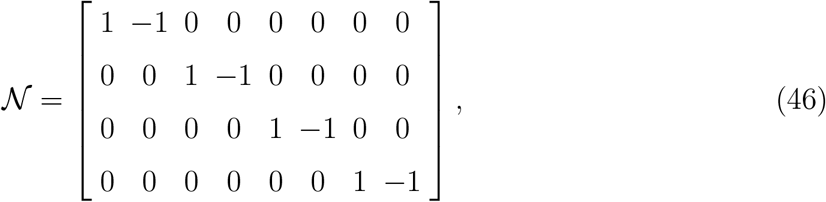

and the reaction rate vector **v**, the diffusion and Jacobian matrices can be obtained from

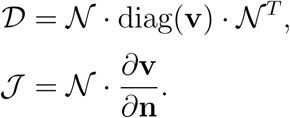

To solve for the correlation matrix 𝒞, we use an algorithm based on the Sylvester equation (ℐ ⊗ 𝒥 + 𝒥^*T*^ ⊗ ℐ) vec(𝒞) = vec(−𝒟), which gives

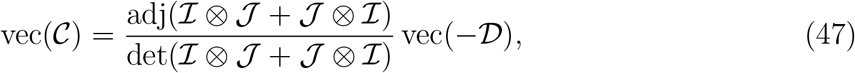

where adj denotes the adjugate matrix, det is the determinant, and ℐ is the identity matrix. The Kronecker product ⊗ is used to find the matrix direct product. That is, for 𝒜 an *m* × *n* matrix and ℬ a *p* × *q* matrix, their Kronecker product ℰ = 𝒜 ⊗ ℬ is a *mp* × *nq* matrix that has elements defined by *e*_*αβ*_ = *a*_*ij*_*b*_*kl*_, where *α* = *p*_*i*−1_ + *k* and *β* = *q*_*j*−1_ + 1. After solving the Sylvester equation, vec (𝒞) can be reshaped to the same dimensions as the Jacobian matrix, leading to the correlation matrix 𝒞, which in turn gives the variance *σ*_*ii*_ and covariance *σ*_*ij*_.

### G. Average noise reduction coefficient

To assess the contribution of each controller modality to reduction of system noise, we define the quantity “average noise reduction coefficient,”denoted as ⟨ *C*_*n*↓_⟩. The controller dimensions include the RFP mean expression, controller strength, controller type (local, global, NCR), controller subtype (MIX, PIX, MIL, PIL), and controller application system (RC or OR). Given a function *ϕ* (*P*_2_, *S*_*c*_, *i, j, k*) that returns noise for a given RFP mean (*P*_2_), controller strength (*S*_*c*_), and discrete dimensions of controller type (*i*), controller subtype (*j*), and controller application case (*k*), we define ⟨ *C*_*n*↓_⟩ of a certain con-troller modality as the average reduction in noise summed across all dimensions, holding the modality constant. We perform this operation after normalizing *ϕ* to the base case through *ϕ*_normalized_ = *ϕ*_experimental case_*/ϕ*_base case_. These considerations lead to

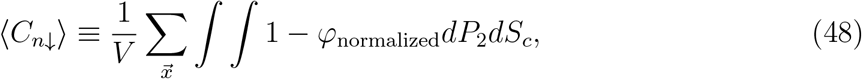

where 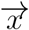 represents all discrete dimensions being summed over and *V* represents the size of the *n*-dimensional region under *ϕ*_normalized_. The coefficient ⟨*C*_*n*↓_⟩ can be used to visualize the average contribution of any controller modality and succinctly compactify a large amount of data into a single number.

### H. Base parameters and parameter rescaling

The resource competition (RC) case is used as the base case. The following parameter values are used unless otherwise specified: 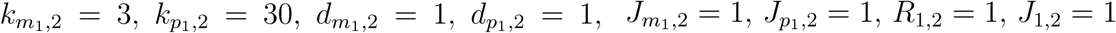, and Ω = 4. The values of *J*_*p*_ and *S*_*c*_ range from 0 to 50. In cases where copy numbers or mean expression levels vary, the values of *R* are in the same range.

Parameter rescaling is done to ensure that the means of the mRNAs and proteins are the same across all cases, which is necessary for fair comparison of the noise behaviors of in all the models. All rescaling is done by resetting the transcription and translation rate constants (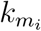 and 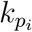) using the resource competition case as the base. The specific values of these rate constants for different systems are listed, as follows.

1. Unlimited Resources (UR) system:

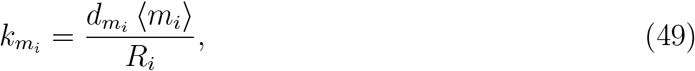

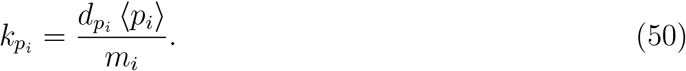
2. Resource Competition (RC) system:

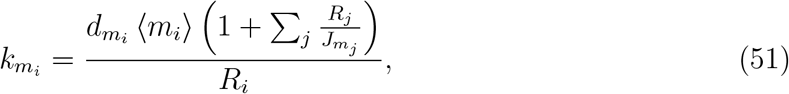

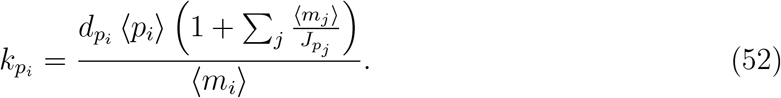
3. Orthogonal Resource (OR) system:

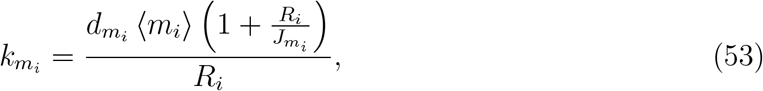

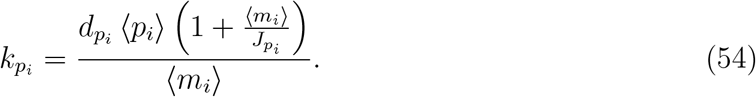
4. RC system with a controller:

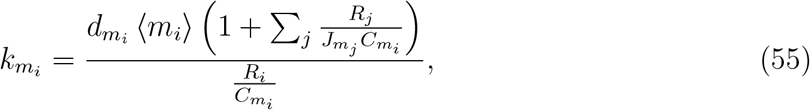

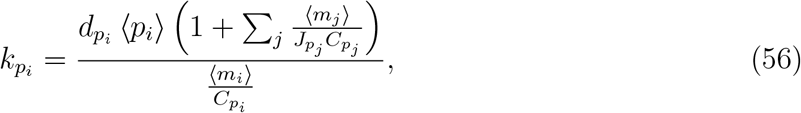

where 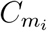 and 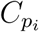 are defined in Eqs. (38) and (39), respectively.
5. OR system with a controller:

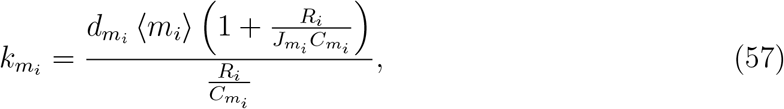

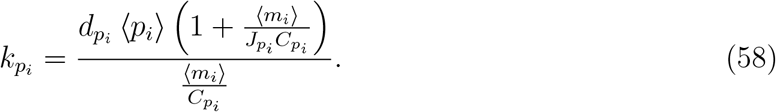

**Supplementary Fig. 1.**
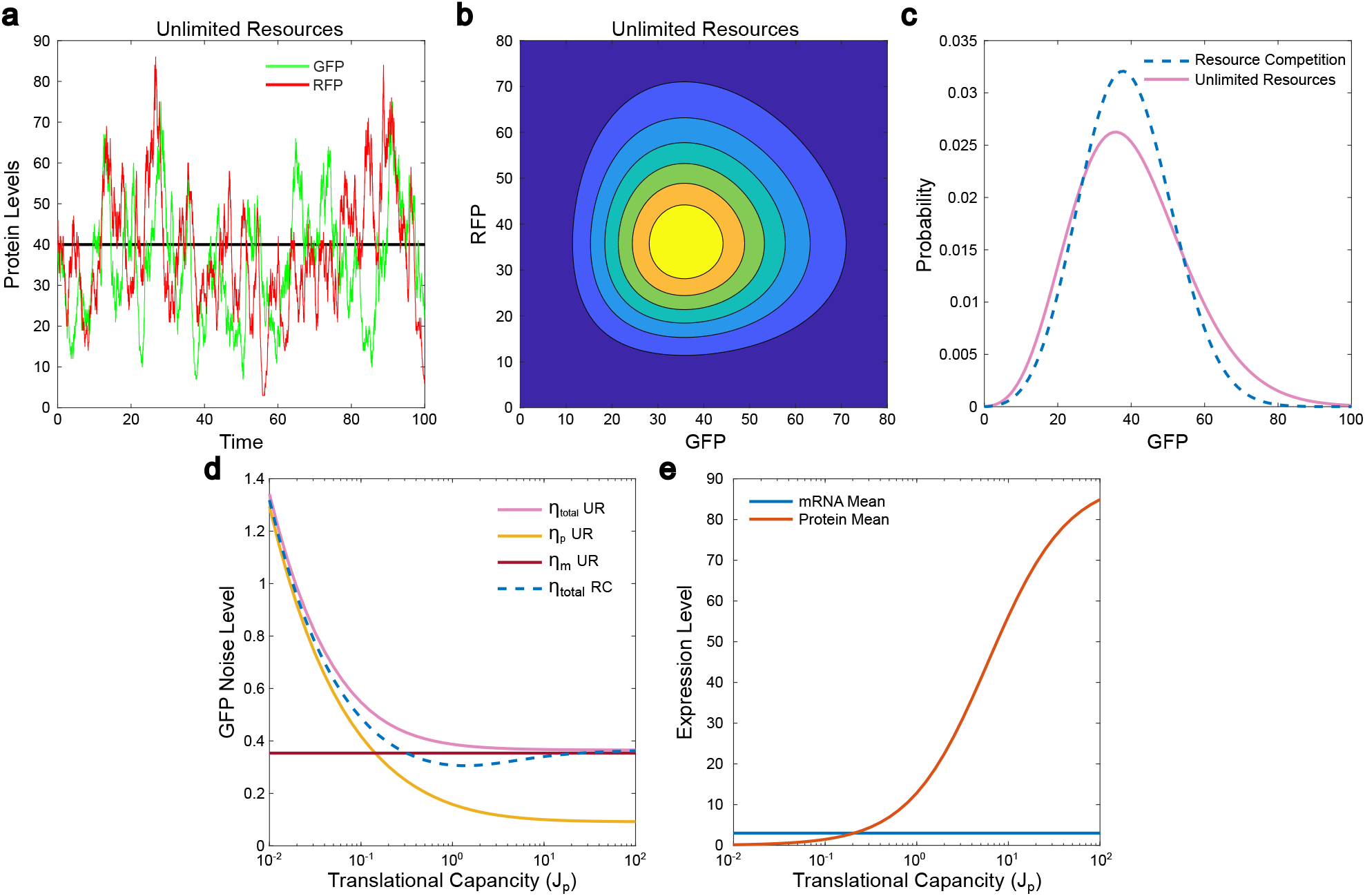
Reduction of gene expression noise by resource competition in comparison with the case of unlimited resources. (a) Gillespie stochastic trajectories of GFP (green trace) and RFP (red trace) expression in a two-gene circuit with unlimited resources. (b) The corresponding distribution of GFP and RFP expression levels from the solutions of the master equation. (c) Distribution of GFP expression levels in the systems with resource competition (blue dashed curve) and unlimited resources (purple curve). (d) The dependence of the GFP total noise level and its decomposition on the translational capacity *J*_*p*_ of limited resources in the host cell for the synthetic gene circuit. The GRP total noise level in the case with resource competition is shown in the dashed curve for comparison. (e) The dependence of mRNA and protein means on the translational capacity *J*_*p*_ for the RC system. The transcription and translation rate constants in the UR model are rescaled to ensure the same means of the mRNAs and proteins in the two models.

**Supplementary Fig. 2.**
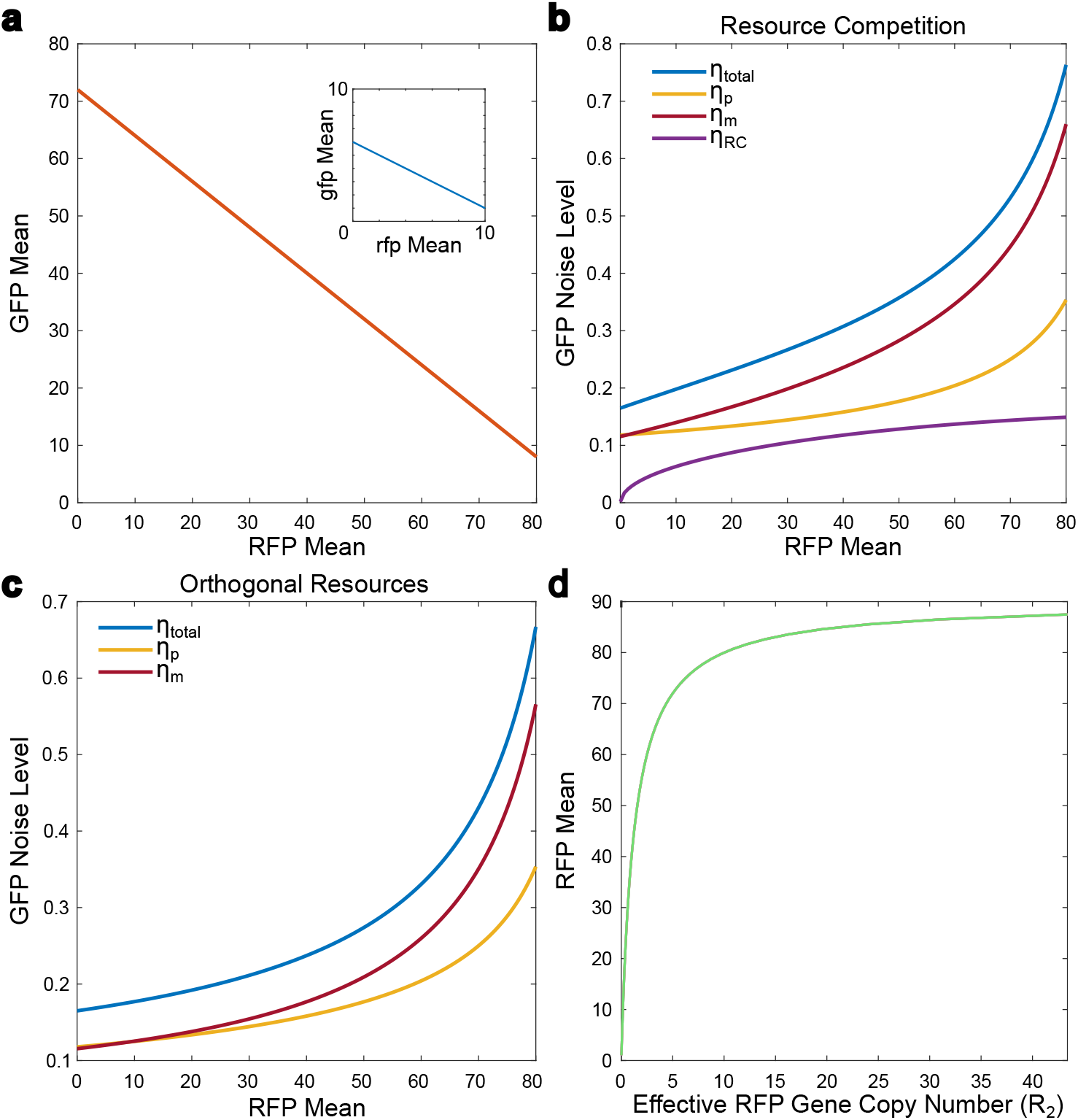
Analysis of protein noise decomposition in the system with and without orthogonal resources. (a) Isoline of GFP mean versus RFP mean in the two-gene system with resource competition. The inset shows the isoline of gfp mRNA versus rfp mRNA, where the effective rfp gene copy number increases. (b) The dependence of the GFP total noise level and its three decompositions on RFP mean in the RC system. (c) The dependence of the GFP total noise level and its two decompositions on RFP mean in the OR system. (d) The dependence of the RFP means on effective rfp copy number.

**Supplementary Fig. 3.**
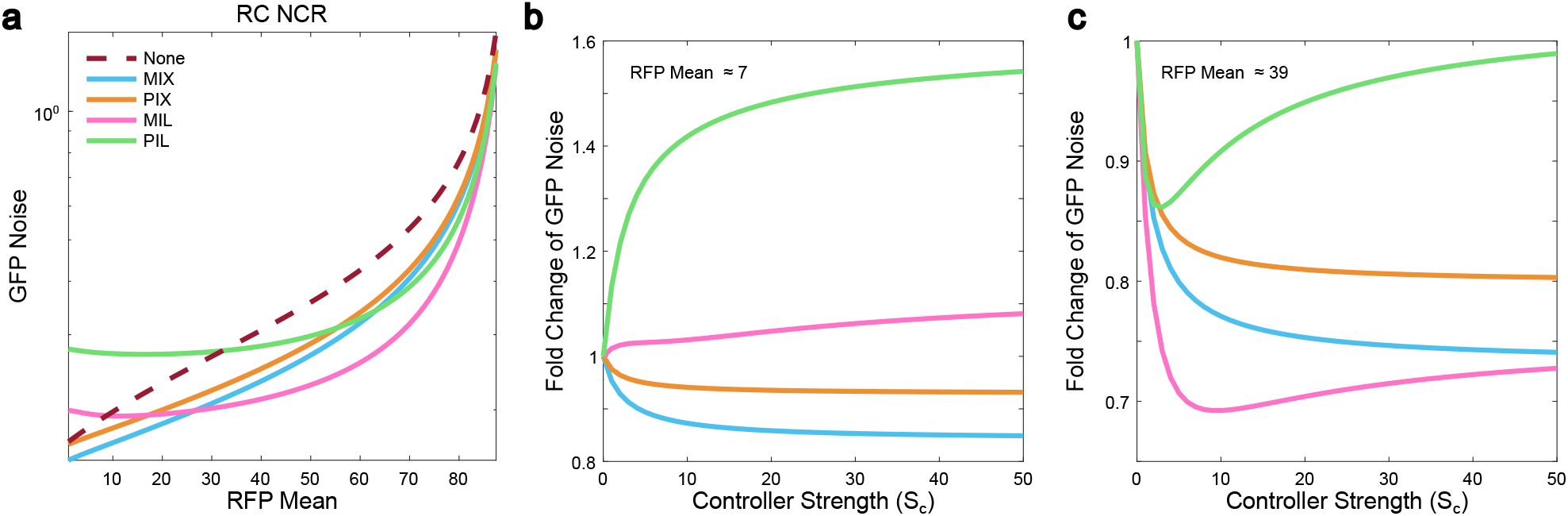
Analysis of the noise reduction by the NCR controllers. (a) Dependence of the non-normalized GFP noise levels on RFP mean for an RC system with no controller (maroon dashed curve), an NCR-MIX controller (light blue curve), an NCR¬-PIX controller (orange curve), an NCR-MIL controller (fuchsia curve), and an NCR-PIL controller (light green curve). (b, c) Dependence of normalized GFP total noise on controller strength with respect to the RC system with no controller with a fixed RFP mean at 6.8 and 38.9, respectively.

**Supplementary Fig. 4.**
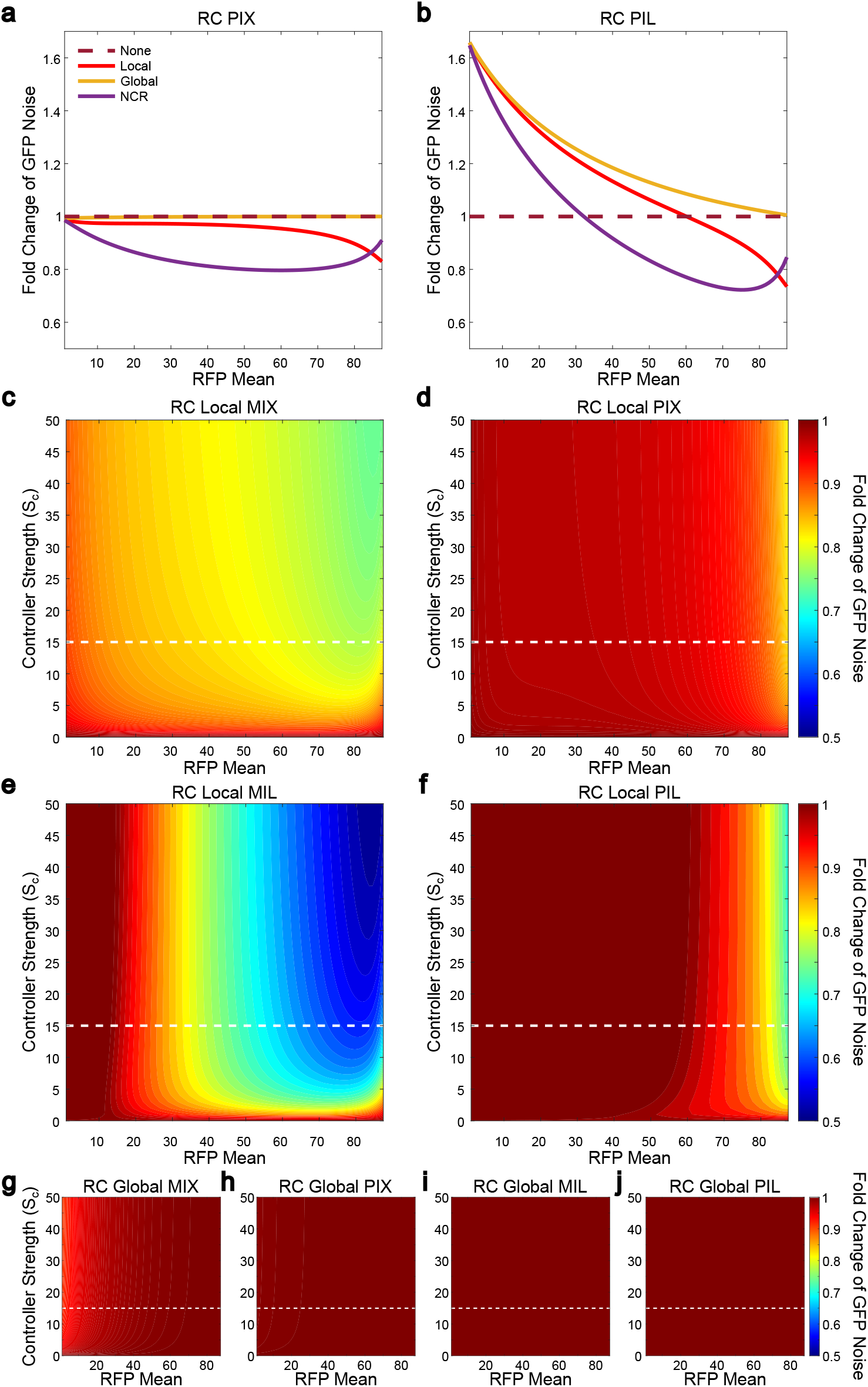
Analysis of noise reduction capability of global and local controllers. (a, b) Dependence of the normalized GFP noise levels on RFP mean for an RC system with, respectively, PIX and PIL subtypes of three controllers. (c-j) Normalized GFP noise levels in the phase plane of RFP mean and controller strength for four subtypes of local controllers (c-f) or global controller (g-j). Deep blue region represents a strong decrease in noise levels in comparison to a resource competition case with no controller, and a deep red region represents a noise level increase or the absence of a noise change. The horizontal white dashed lines represent the controller strength Figs. 4c, 4d, 3a, and 3b.

**Supplementary Fig. 5.**
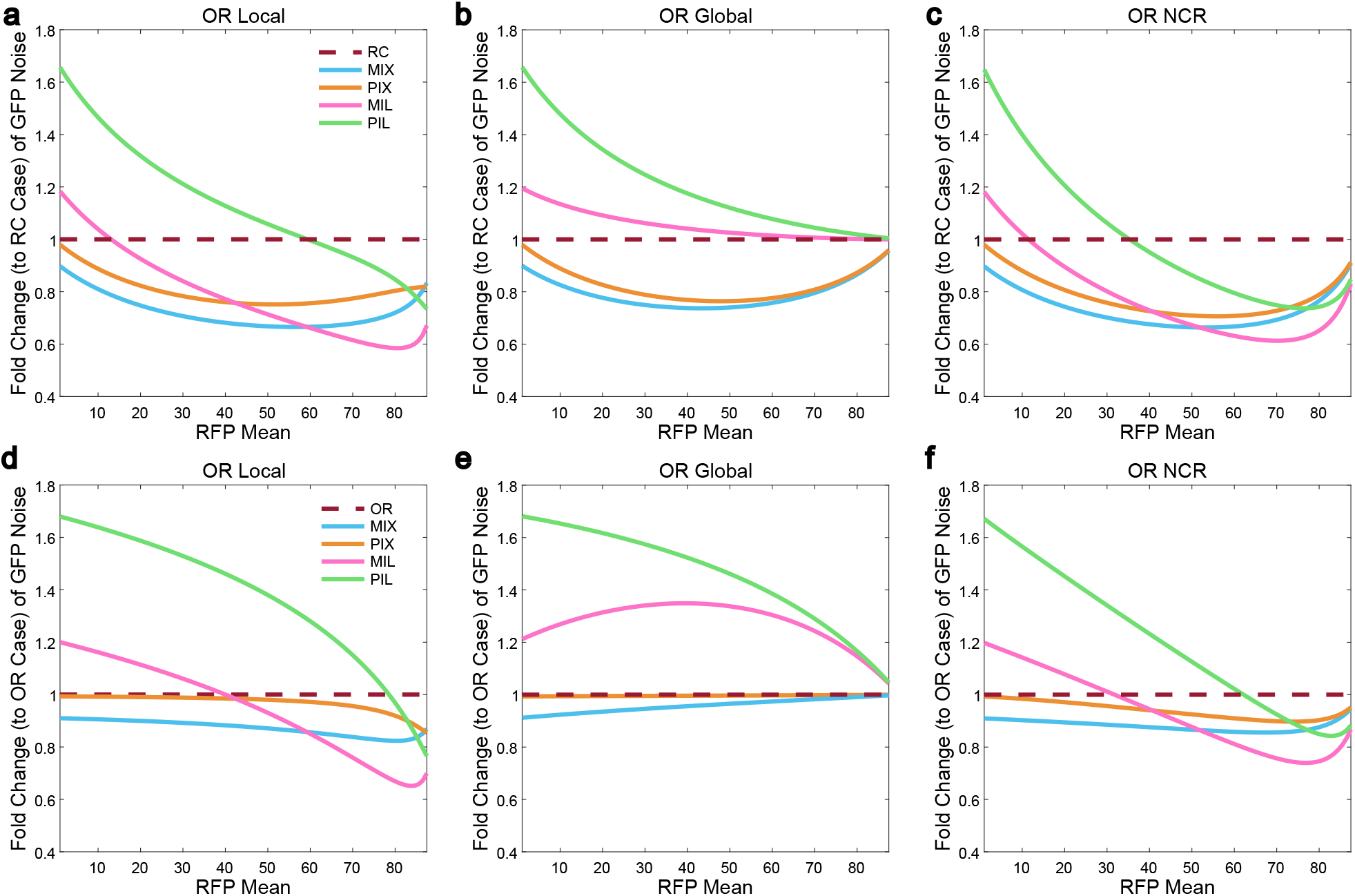
Analysis of the noise reduction by the negative feedback controllers combined with orthogonal resources. (a-c) Dependence of the normalized GFP noise levels on RFP mean for a system with, respectively, four subtypes of local controller, a global controller, and an NCR controller. The GFP noise levels are normalized to the case with no controller and no orthogonal resources. (d-f) Same as a-b, but with the noise levels normalized to the case with orthogonal resources.

